# Aberrant iron deposition in the multiple sclerosis spinal cord relates to neurodegeneration

**DOI:** 10.1101/2024.10.25.619794

**Authors:** Marco Pisa, Andrew Lockhart, Thomas Angell, Aimee Avery, Alex Waldman, Carla Rodriguez, Tiago Duarte, Jonathan Pansieri, Zaenab Dhari, Mary Bailey, Simon Hametner, Hal Drakesmith, Monika Hofer, Clara Limbaeck, Gabriele C DeLuca

**Affiliations:** Nuffield Department of Clinical Neurosciences, University of Oxford; REQUIMTE-LAQV, Departamento de Química, Faculdade de Ciências e Tecnologia, Universidade Nova de Lisboa; i3S-Instituto de Investigação e Inovação em Saúde, Universidade do Porto; Mandell MS Center, Trinity Health of New England; Department of neuroimmunology, Medical University of Vienna; MRC Translational Immune Discovery Unit, MRC Weatherall Institute of Molecular Medicine, Radcliffe Department of Medicine, University of Oxford, Oxford, UK; Neuropathology, Oxford University Hospital NHS Fundation Trust

## Abstract

**Background:** Iron accumulates in microglia-macrophages at the edge of multiple sclerosis (MS) lesions in the brain. Iron-rimmed brain lesions strongly predict disability accumulation, supporting iron metabolism is crucial in MS pathology. Little is known about iron distribution in the spinal cord.

**Methods:** Autopsy cervical, thoracic and lumbar spinal cord samples from 9 controls and 46 MS donors of whom a subset (n=36) had mesiofrontal motor cortical tissue available for study, were labelled and systematically assessed for iron (DAB-enhanced Turnbull), myelin (PLP), axons (Palmgren silver), microglia-macrophages (TMEM119, Iba1, CD68), astroglia (GFAP), oligodendroglia (OLIG2), acute axonal injury (B-APP, SMI-32, NPY-1R) and oxidative stress (E06). MS lesional and non-lesional areas were considered. Total non-haem iron was quantified by inductively coupled plasma optical emission spectroscopy (ICP-OES).

**Results:** In controls, iron predominantly localised to oligodendrocytes with total non-haem iron relating to total myelin fraction, which markedly differed in MS where iron accumulated in microglia-macrophages, subpial astrocytes, and axons in non-lesional areas. Iron laden microglia-macrophages were over-represented relative to total microglial-macrophages and displayed dysmorphic features. Iron-positive axons showed a disto-proximal gradient (highest at lumbar level) with a predilection for the corticospinal tracts. The extent of iron axon positivity related to smaller spinal cord area, lower total axonal counts, and greater oxidative stress. Iron positivity in each cellular compartment (i.e. subpial astrocytes, microglia-macrophage and axons) related to one-another and total non-haem iron correlated with axonal counts in MS. No iron-rimmed lesions were detected in the spinal cord unlike in cortical grey and subcortical white matter of the same cases where 22% and 80% of iron-rimmed lesions, respectively, were seen.

**Conclusions:** Despite the conspicuous absence of iron-rimmed lesions in the MS spinal cord, we demonstrate widespread aberrant iron distribution in the MS spinal cord that relates to oxidative stress and neurodegeneration independent of demyelination. The distal cord predominant and corticospinal tract specific accumulation of iron in axons mirrors the pattern of length-dependent motoric disability commonly encountered in progressive MS. These findings implicate aberrant iron accumulation as a novel, clinically relevant, feature of MS spinal cord pathology.

## 1. Introduction

The traditional hallmark of multiple sclerosis (MS) pathology is the presence of focal inflammatory demyelinating lesions forming around post-capillary venules and veins.^1^ Clinically, lesion development associates with the emergence of reversible focal neurological symptoms (i.e. relapses), is well captured by standard MRI imaging techniques, and effectively prevented by currently available disease modifying therapies.^2^ However, most people with MS eventually enter a progressive phase, in which accumulation of irreversible disability and diffuse neuroaxonal loss occur in an age-dependent manner, independent of new lesion formation or exposure to effective disease modifying therapies.^3–6^ Contrary to the heterogeneous clinical presentations of relapsing MS, progressive MS is characterised by a motor-predominant myelopathy with a length-dependent gradient of severity, for reasons that are unknown.^4,5^ Widespread chronic inflammation, leading to oxidative stress, mitochondrial failure, and accelerated senescence are thought important contributors.^7,8^

In recent years, the discovery that paramagnetic-rim lesions (PRLs), which identify iron-rich microglia-macrophages at the edge of brain lesions, associate with markers of neurodegeneration and irreversible disability, has shifted focus to the role of iron in processes relevant to progressive MS.^9,10^ However, the presence of PRLs in the spinal cord, arguably the most clinically relevant for disability progression in MS, is debatable from an imaging standpoint.^11^ Pathological, imaging, and genetic studies support a role for iron and its metabolism as a driver of disability progression.^12–17^ For example, increased iron concentration in the deep grey matter (GM) has been associated with oxidative burst and increased neurodegeneration and atrophy,^13^ while reduced iron content in non-lesional white matter (WM) has been observed with increasing disease duration.^12^ Most studies have focused on the characterisation of iron in the lesional and perilesional areas in the brain. Little is known about iron metabolism in the spinal cord and there has been no systematic characterisation of iron in non-lesional tissue reported to date.

We hypothesised that aberrant iron deposition in the spinal cord contributes to neurodegenerative outcomes and disability in MS. Thus, we sought to characterise the distribution and extent of iron in the MS spinal cord tissue and its relation to pathological and disability outcomes.

In a large autopsy cohort of progressive MS cases, we note the conspicuous absence of iron-rimmed lesions in the MS spinal cord unlike in the brain of the same cases where iron-rimmed lesions were found at a frequency as previously described. Interestingly, we describe aberrant iron accumulation in axons, microglia-macrophages, and subpial astrocytes of the non-lesional spinal cord. We observe a clearly length-dependent and motor tract predominant pattern of iron deposition in MS, which corresponds to the clinical phenotype commonly encountered in progressive MS. In support of a disability-relevant role, iron accumulation related to increased oxidative stress, reduced axonal count, and greater cord atrophy. These findings implicate aberrant iron accumulation as a novel, clinically relevant, feature of MS spinal cord pathology.

## 2. Materials and Methods

### 2.1 Study cohort

A well-defined post-mortem cohort of pathologically confirmed PMS cases (n=46) and non-neurological controls (n=8) from the UK MS Tissue Bank were used. All cases had cervical, thoracic, and lumbar cord tissue for study whilst mesiofrontal motor cortical samples were available for 36 MS cases. All tissue was used with appropriate ethical approval as per the Human Tissue Act 2004 (Imperial-Research Ethics Committee (REC) approval number: 08/MRE09/31+5).

### 2.2 Experimental procedure

Formalin-fixed paraffin-embedded 6µm-thick sections from each case and level of the neuraxis were labelled for non-haem iron using an optimised Turnbull protocol [Diaminobenzidine (DAB)-Enhanced Turnbull Staining Protocol (Tirmann-Schmeltzer Method)].^12,18^ The specificity of the Turnbull reaction was confirmed by staining sections after 72h incubation in either a 0.1M citrate pH1.0 or in a 0.1M EDTA buffer, which both abolished the Turnbull signal, as previously reported (**Supplementary Data 1**).^19^

Adjacent sections were immunolabelled for myelin (PLP, BioRad, #MCA839G), microglia-macrophages (Iba-1, Wako, #019-19741 and CD68, Dako, #M087601-2), oligodendroglia (Olig-2, IBL-America, #18953), astrocytes (GFAP, Dako, #Z0334), oxidative stress (E06, Absolute Ab, #Ab02746-1.1), and acute axonal injury (BAPP, Invitrogen #13-0200; SMI-32, Merck Millipore, NF-H Non-phosphorylated #Ne1023; NPY-1R, Origene, #TA346235) (**Supplementary Data 2**). Spinal cord sections were impregnated with Palmgren’s silver to demonstrate axons, as previously published.^20,21^

A subset of cases were double-labelled for Turnbull and the above listed oligodendroglial, microglia-macrophage and astrocytes markers, and acute axonal injury. For double labelling, sections underwent DAB-enhanced Turnbull staining and subsequent immunolabelling using the same above described methods, except that immunolabelling was developed using Vector Red (Vectastain ABC-AP kit).

### 2.3 Neuropathological assessment

Qualitative evaluation was conducted by four assessors (MP, AL, CL, GDL; **Figure 1**) and subsequently systematically recorded by two independent assessors (MP and AL). Based on the characteristic reactivity patterns, a tailored semiquantitative scoring system of spinal cord iron distribution was designed, which two assessors (MP and AL) then used to assign scores independently. In cases of disagreement between assessors, slides were re-reviewed and a consensus score assigned.

**Figure 1:**
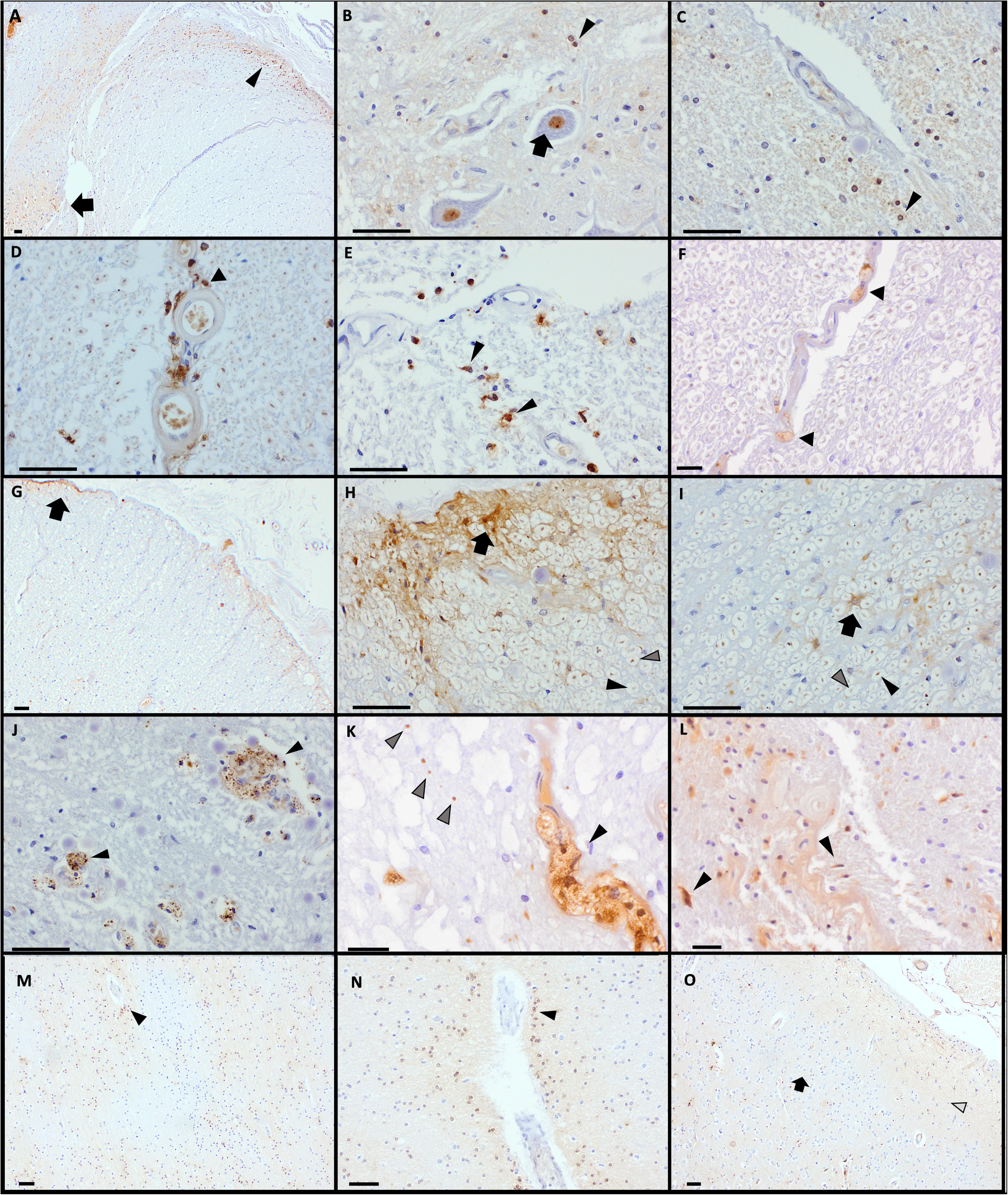
Characteristics of iron distribution in the the spinal cord (**A-F**) and motor cortex (**M-O**); all sections are stained using DAB-enhanced Turnbull. In controls, most non-heme iron was contained in oligodendrocytes(**A-C**). Clusters of iron-positive oligodendrocytes were most frequently identified in the dorsal entry root area (**A**), the subpial area (**A,** arrowhead) and in the grey matter dorsal horn (**A**, arrow), at the grey-white matter junction (**B**, arrowhead), and in close proximity to white and grey matter vessels (**C,** arrowhead). Ventral horn neurons frequently showed an intense nuclear and nucleolar staining pattern (**C**, arrow). Iron-positive microglia-macrophages were frequently found in the non-lesional white matter in MS cases (**D-F**). Infiltrating iron-positive microglia-macrophages were common in the non-lesional white matter of MS cases, typically in proximity to a vessel (**E-F**) and frequently displayed a dystrophic morphology (**F**, arrowhead). Rarely, intravascular foamy macrophages with a distinct iron-positive granular cytoplasmic staining were also observed (**D**). Hypertrophic astrocytes with thick iron-rich processes were observed in the subpial region of MS cases (**G,** arrow & **H,** arrow) and rarely in non-lesional white matter where it tended to associate with iron-rich axons (**I**, arrow). Sparse iron-positive axons (**H-I,** black arrowhead), with adjacent iron-negative axons (**H-I**, grey arrowhead) were found in close proximity to iron-positive hypertrophic astrocytes in both the subpial area (**H**) and in non-lesional tissue (**I**). In plaques, the most striking finding was the presence of foamy macrophages with iron-rich cytoplasmic inclusions and often forming rich perivascular cuffs (**J-K**, black arrowheads). In active plaques, iron-positive axonal swelling (**K,** grey arrowheads), a blush of extracellular iron (**L**), and sparse iron-positive glial cells with astrocytic morphology (**L**, black arrowheads) were occasionally found. In the mesiofrontal motor cortex, a more diffuse and intense Turnbull reactivity is detected in the subpial white (**M-N**) and grey matter (**O**) compared to the spinal cord. In the white matter, iron-positive oligodendrocytes clustered around vessels (**M-N**, black arrowheads) generating a tigroid pattern in the tissue (**M**). In the grey matter, layer II was particularly devoid of iron (**O**, arrow) while the subpial layer I showed a variable degree of fibrillar iron-positive staining (**O**, grey arrowhead). Scale bar corresponds to 20µm.

#### Characterisation of non-haem iron in the non-lesional spinal cord

For each case, iron-positive axons, total glia, and microglia-macrophages were quantified in each side of the following spinal cord WM areas (i.e. anterior, antero-lateral, postero-lateral and posterior) and in GM using a semiquantitative scoring system. Assessors were blinded for case demographic and pathological characteristics. Tracts containing demyelination were excluded to derive average non-lesional spinal cord measures for downstream analyses.

1. Iron-positive axonal score was based on the observation that iron-positive axons tended to occur in clusters: 0, no clusters; 1, cluster of iron-positive axons confined to a 200X field of view; 2, cluster spreading beyond a 200X field of view
2. Iron-positive total glial score was defined by the number of iron-positive glial cells in the 200X field of view with the highest density: 0, none; 1, ≤5 cells; 2 >5 cells. A similar score was given only considering the cells with typical activated microglia-macrophage morphology (foamy, ameboid or dysmorphic)
3. Subpial spinal cord iron-positive reactivity was quantified based on extent perimeter coverage: 0, none; 1, present around ≤ 1/3 of the cord perimeter; 2, between 1/3 and 2/3; 3 if > 2/3 of cord perimeter
4. In spinal cord GM, the extent of iron-positive glial cells was recorded as: 0, absent; 1, iron-positive cells only present in the dorsal horn; 2, iron-positive glia present across the whole GM area (i.e. dorsal and ventral horns)
5. Iron-positive neuronal score was based on the proportion of ventral horn motor neurons with visible nucleoli displaying Turnbull reactivity: 0, none; 1, < 50%, 2, ≥ 50%

#### Quantitative analysis of non-haem iron distribution in the glial compartment in the non-lesional spinal cord

A subset of MS cases (n=10) and controls (n=5) propensity-score matched for age at death, sex, and total iron-positive glia score were selected for additional quantitative analyses. Iba-1 and Olig-2 expression were quantified in selected 40X magnification fields of view (FOVs) in pre-defined trajectories, selected for quantification in non-lesional areas of notional tracts (anterior corticospinal tracts (n=6), lateral corticospinal tracts (n=10), and dorsal columns (n=12) (**Supplementary Data 3**), as previously described.^20,21^ Olig-2 positive cells were quantified using an optimised positive cell count macro on QuPath,^22^ whilst a positive pixel count macro was used for Iba-1 expression given the non-nuclear expression pattern. In the same FOVs of adjacent double-labelled (i.e. Iba-1 plus Turnbull, and Olig-2 plus Turnbull) sections, the number of iron-positive microglia-macrophages and oligodendrocytes were manually counted by two independent assessors after blinding case identifier on each section (MP & AL, and MP & TA, respectively) (**Supplementary Data 3**), and related to total number of cells.

#### Association of iron-positive axons with markers of axonal injury and oxidative stress in the non-lesional spinal cord

A subset of MS cases lying on the extremes of iron-positive axon clusters were further evaluated. To accomplish this, 10 MS cases and 5 controls propensity-score matched for age at death and sex were selected. MS cases were selected so that 5 had an average axonal score of 0 (i.e. 5 with and 5 without iron-positive axonal clusters). The total number of iron-positive axons were manually counted in each FOV by an assessor (TA) blinded to case identifier and verified by an independent assessor (MP). Optimised positive element count macros were developed on QuPath to quantify axons positive for E06, B-APP, SMI-32, and NPY-1R (**Supplementary Data 3**).

#### Quantification of myelin fraction, microglial inflammation and axons in the non-lesional spinal cord

Myelin fraction was calculated as the ratio between the PLP-positive area and the spinal cord section area. The PLP-positive area was measured using a high resolution (0.5 µm/pixel) positive-pixel threshold macro on QuPath to allow the identification of area covered by myelin wraps in each section (**Supplementary Data 4**).

PLP-stained sections were used to guide delineation of lesional GM and WM areas so that microglia-macrophage inflammation in non-lesional areas could be measured. The expression of the pan-marker, Iba-1, was quantified in the region of interest using an optimised positive pixel count macro on QuPath.

Palmgren’s silver stained sections were used for axonal count in non-lesional FOVs in the anterior and lateral-corticospinal tracts and in the dorsal column. Total and notional tract (i.e. dorsal column, lateral corticospinal tract, and anterior corticospinal tract) cross-sectional spinal cord areas and total axonal number estimates were obtained, as previously published.^20,21^

#### Characterisation of demyelinating plaques in the spinal cord and motor cortex

PLP- and CD68-stained sections were used to classify WM plaques into active, mixed active-inactive, and chronic inactive based on established criteria.^23^ Cortical lesions were classified into type I-IV, as previously described.^5^ For each demyelinated lesion, the presence of an iron-rim at the plaque edge was systematically noted. Presence of iron-positive microglia-macrophages was scored (0, none; 1, few perivascular foamy macrophages; 2, full macrophage perivascular rim).

### 2.4 Quantification of non-haem iron in the spinal cord using ICP-OES

Two 15µm FFPE sections were deparaffinised and incubated in a 1mL 0.1M sodium citrate solution pH 1.0 for 72h. The buffer was extracted, and iron was quantified using ICP-OES (**Supplementary Data 5**). The inductively coupled plasma optical emission spectroscopy (ICP-OES) analysis was carried out on a sequential ICP (Ultima, Horiba Jobin Yvon, France), using the Horiba Jobin Yvon ICP Analyst software (v. 5.4) as detailed in **Supplementary Data 5**. Iron concentration in the buffer was normalised by tissue volume (spinal cord area in µm^2^ * 30µm) and log10 transformed for analyses.

### 2.5 Statistical analyses

All analyses were carried out using IBM SPSS v27.0. Generalised estimating equations were used to compare neuropathological variables in each sample (iron semiquantitative scores, glial and axonal markers, and ICP-OES iron concentration values) between MS and controls, or between MS cases with and without presence of iron-positive axonal clusters using a correlation matrix representing the within-subject dependencies. Cord level and myelin fraction were included as covariate, when appropriate. Comparison of the frequency of iron features between groups were analysed with cross-tabs; Pearson χ2 or linear-by-linear correlation p-values are reported for categorical and ordinal values, respectively. Marginal estimated mean and CI95% are reported for each group, together with the Wald χ2 and Bonferroni adjusted p-values of the pairwise comparisons. Correlations between iron semiquantitative scores, glial and axonal markers, and ICP-OES iron concentration values were assessed by fitting linear generalised estimating equation models; models were corrected for cord level (ordinal variable) and myelin fraction, when appropriate. Wald χ2 and p-value of the independent variable are reported for significant models. For graphical purposes, comparison of medians between groups were run in addition to the GEE models described above, using Mann-Whitney (for binary categories) or Kruskal Wallis tests with post-hoc Tukey test (if >2 classes); comparison of medians for paired values (i.e. iron scores between funiculi of the same section) were analysed with Friedmann’s tests. The threshold for statistical significance was set at p<0.05 (* indicates p<0.05, ** indicates p<0.01).

## 3. Results

### 3.1 Population characteristics

Cohort characteristics of the 46 progressive MS and 8 non-neurological cases studied are reported in **Table 1**. Cervical, thoracic, and lumbar cord sections were included when available, for a total of 136 MS and 22 control spinal cord blocks studied. Controls showed older age at death and longer post-mortem interval compared to the MS cohort (both p=0.03). In a subset of 36 MS cases, mesiofrontal motor cortical samples were available and have been included in the study. No difference in demographic characteristics was observed between the motor cortical and the spinal cord cohorts.

**Table 1.**
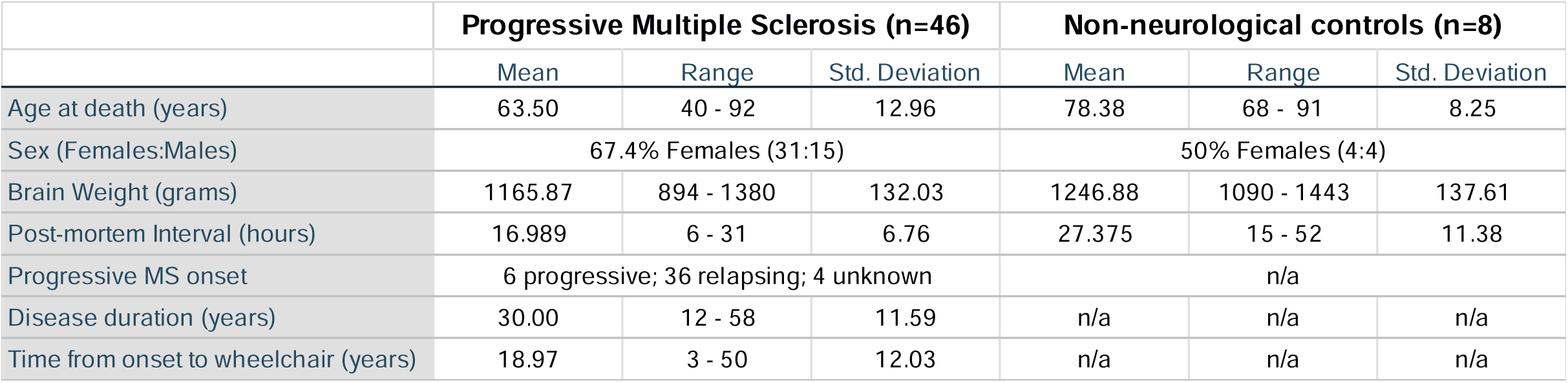
Population characteristics.

|  | Progressive Multiple Sclerosis (n=46) |  |  | Non-neurological controls (n=8) |  |  |
| --- | --- | --- | --- | --- | --- | --- |
|  | Mean | Range | Std. Deviation | Mean | Range | Std. Deviation |
| Age at death (years) | 63.50 | 40 - 92 | 12.96 | 78.38 | 68 - 91 | 8.25 |
| Sex (Females:Males) | 67.4% Females (31:15) |  |  | 50% Females (4:4) |  |  |
| Brain Weight (grams) | 1165.87 | 894 - 1380 | 132.03 | 1246.88 | 1090 - 1443 | 137.61 |
| Post-mortem Interval (hours) | 16.989 | 6 - 31 | 6.76 | 27.375 | 15 - 52 | 11.38 |
| Progressive MS onset | 6 progressive; 36 relapsing; 4 unknown |  |  | n/a |  |  |
| Disease duration (years) | 30.00 | 12 - 58 | 11.59 | n/a | n/a | n/a |
| Time from onset to wheelchair (years) | 18.97 | 3 - 50 | 12.03 | n/a | n/a | n/a |

### 3.2 Iron distribution in the non-lesional spinal cord

As reported by Morris and colleagues in 1992^25^, spinal cord tissue from controls showed little non-haem iron. In controls, Turnbull reactivity was predominantly observed around the nuclei of glial cells with oligodendrocyte morphology (i.e. small round nuclei and a perinuclear halo) which were surrounded by a small cloud of fine iron-positive processes. In a subset of cases, these were confirmed to double label with the oligodendroglial marker, Olig-2. Iron-positive oligodendrocytes were most commonly found in small clusters in the dorsal root entry zone and in adjacent dorsal and lateral column subpial areas (**Figure 1A**). Iron-positive oligodendrocytes were also found in close proximity to WM and GM vessels, at the GM-WM junction (**Figure 1B**), and in the dorsal horn. Perinuclear and nucleolar iron reactivity in ventral horn neurons was observed in 42% of cases (**Figure 1B**) (no difference between MS and controls, p>0.3), consistent with previous descriptions.^26^ Finally, in the subpial region, astrocyte processes showed varying degrees of iron-positivity.

In contrast, MS cases exhibited greater variability in Turnbull staining between cases, with some features being rarely observed in controls and specifically targeted for systematic quantification. First, MS cases displayed a marked increase in iron-positive axons compared to controls, particularly within motor pathways (i.e., anterior and lateral corticospinal tracts) where a disto-proximal gradient of staining was observed. Second, iron-positive microglia-macrophages were frequently observed infiltrating non-lesional WM (Figure 1E,F). Finally, subpial astrocyte processes were more commonly found in MS cases, often forming a dense fibrillar network delineating the spinal cord perimeter (**Figure1 G,H**).

### Iron-positive axons in non-lesional white matter

In MS cases, large clusters of iron-positive axons were seen, particularly in the lateral cortico-spinal tract area, while only rarely seen in controls. Interestingly, iron-positive axons almost invariably displayed a symmetrical pattern in the two hemi-cord areas of non-lesional tissue.

Iron-positive axons were detected in 43.7% (55/126) of MS and 13.6% (3/22) of control samples (χ2 7.08, p=0.008). In controls, iron-positive axons were sparse when present. In fact, iron-positive axonal score was over 4-fold increased in MS (0.26 CI95% 0.17 – 0.35) compared to controls (0.06 CI95% −0.03 – 0.16; Wald χ2 33.59, p<0.001) (**Figure 2**).

**Figure 2:**
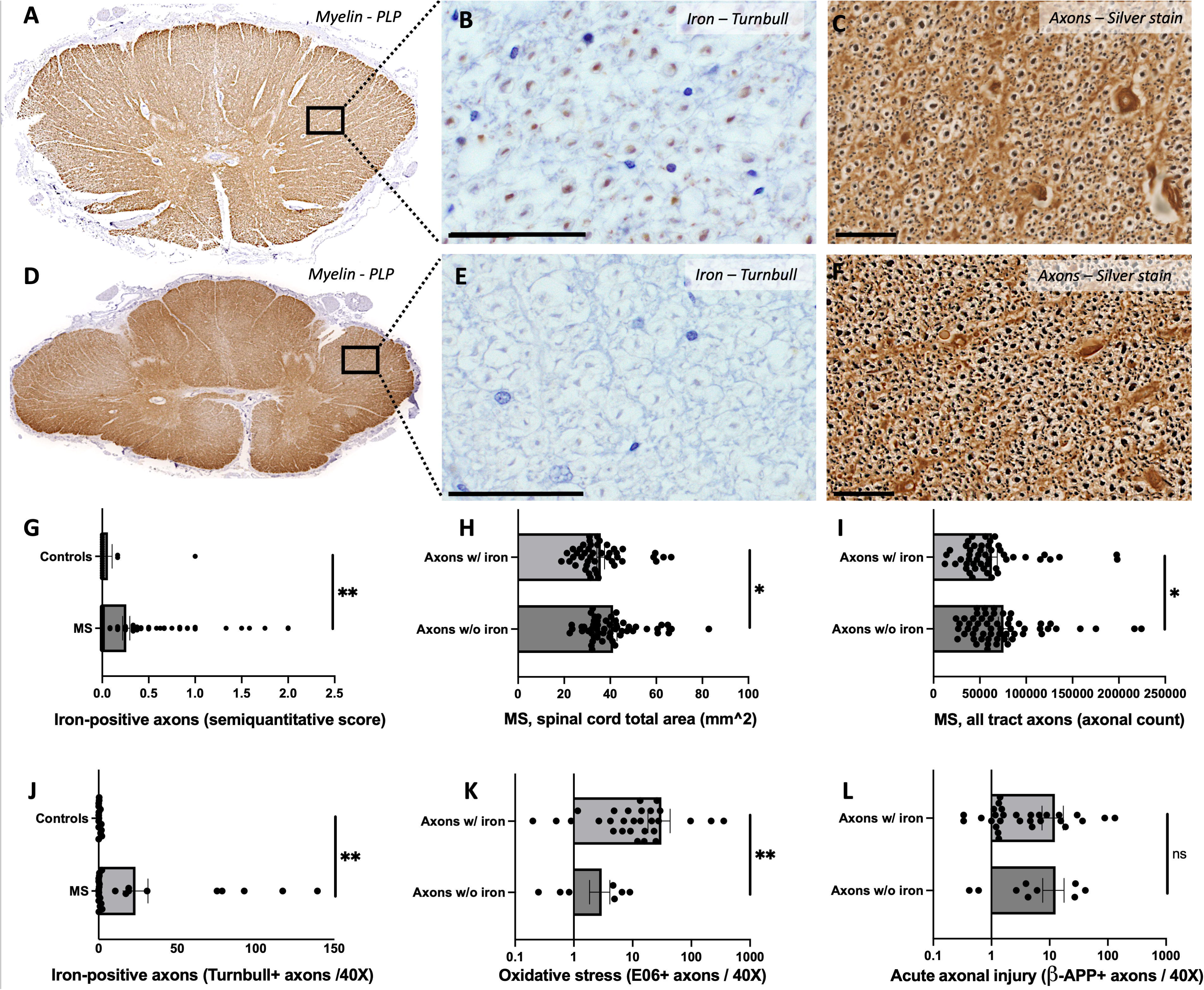
Iron-rich axons are common in the non-lesional white matter and associate with reduced spinal cord area, lower axonal counts and increased oxidative stress. Two spinal cord sections from two progressive MS cases are displayed (A-C, D-F). In non-lesional areas (i.e., normal myelin A, D), non-heme iron was detected in 44.7% of cases (axons with iron in B, axons without iron in E). Cases with iron-rich axons tended to display a symmetrical pattern of Turnbull axonal staining, with predominant involvement of lateral cortico-spinal tracts (LCST) and of the perivascular axons. The presence of this iron-rich axonal pattern associated with lower spinal cord area (G) and axonal counts (H); this was confirmed also considering the LCST alone (I). In a subset of 10 MS and 5 controls cases, manual axonal counting in each spinal cord tract confirmed the increased iron-positive axons in MS (J). Presence of iron-positive axons in a tract associated with higher oxidative stress (E06, oxidated phospholipids; K) but not with markers of acute axonal injury (Beta-amyloid precursor protein; L). Scale bar in B-C and E-F corresponds to 50µm.

Iron-positive axons were more frequently observed in the posterior aspect of the postero-lateral funiculi compared to the anterior, posterior aspect of the lateral funiculi, and dorsal columns in both MS and controls (Friedman’s both p<0.001). When focusing on the iron-positive axons in the lateral funiculus, preferential involvement of the notional lateral cortico-spinal tract (LCST) was found in 54.35% of the cases with entire lateral funicular staining being observed in 32.61% of cases. A small number of cases displayed preferential iron-positivity in axons near the subpial zone (13.04%) (**Figure 3**).

**Figure 3:**
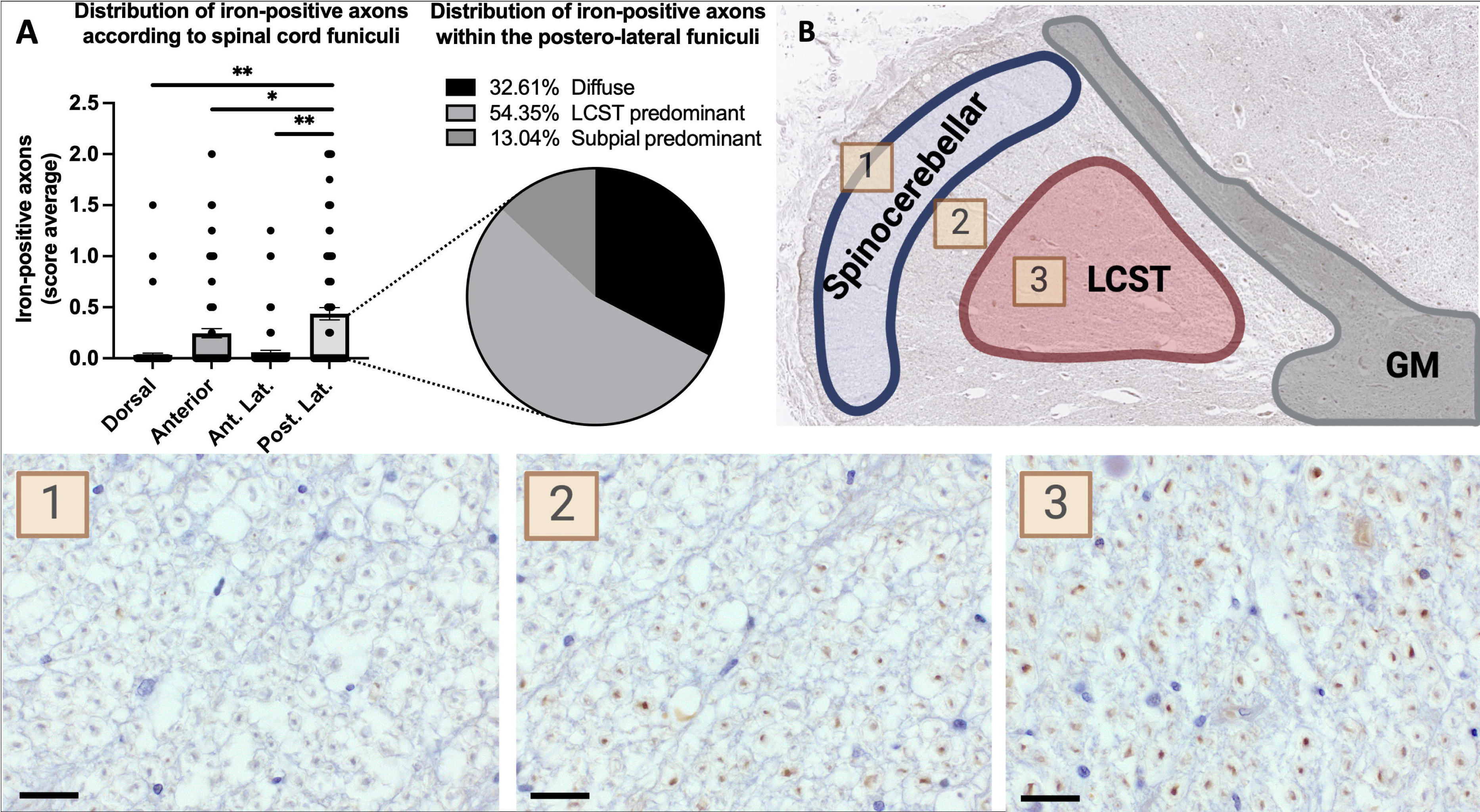
Iron-rich axons display a predilection for corticospinal motor tracts and a disto-proximal gradient of distribution. Distribution of iron-positive axons shows a predilection for the anterior and – even more so – for the postero-lateral tracts (A). When focusing specifically on the distribution of iron-axons in the postero-lateral tract, most cases showed a distinct predominance for the lateral cortico-spinal tract (A). A representative spinal cord section from a progressive MS case stained with DAB-enhanced Turnbull demonstrates a clear selectivity for lateral cortico-spinal tract (B): **Magnification 1** shows only few faintly stained axons in the spinocerebellar tract, **Magnification 2** shows several iron-positive axons in the bottom right side of the picture, **Magnification 3** shows a homogeneous iron-positive staining in the great majority of the lateral corticospinal tract. Scale bar corresponds to 20µm.

A disto-proximal gradient, with a trend towards increasing prevalence and density of iron-positive axons in lumbar compared to cervical cord level, was observed. Specifically, iron-positive axons were not observed in non-lesional motor cortex (0/36, as discussed in section **3.4**) with increasing prevalence of iron-positive axons being found in cervical (14/40, 35%), thoracic (19/42, 45.2%), and lumbar (22/44, 50%) cord levels. Higher semiquantitative iron-positive axon scores were also observed more distally along the cord (cervical: 0.17, CI95% 0.07 – 0.26, thoracic: 0.28, CI95% 0.15 – 0.41, lumbar: 0.32, CI95% 0.18 – 0.48; Wald χ2 4.7, p=0.09).

Higher iron-positive axon score predicted lower total axonal counts (Wald χ2 7.21, p=0.007) and smaller cord areas (Wald χ2 4.88, p=0.027), and in particular iron-positive axons in the LCST predicted total (Wald χ2 5.35, p=0.021) and smaller cord areas (Wald χ2 6.73, p=0.009). All the analyses were confirmed correcting for cord level (**Figure 2**). No correlation was found between iron-positive axons and demographic variables, such as age at death, sex, or post-mortem interval.

In a cohort of 15 cases, manual counting of iron-positive axons confirmed a 40-fold increase in Turnbull-positive axons in MS cases compared to controls (Wald χ2=4.642, p=0.031), as well as predilection for anterior and lateral corticospinal tracts compared to the dorsal columns (Friedman’s p<0.001). We then analysed the association between Turnbull-positive axons and axons positive for the oxidative stress marker, E06 (i.e. oxidated phospholipids), and acute axonal injury. Tracts with a higher number of Turnbull-positive axons exhibited an over five-fold higher number of E06-positive axons (Wald chi-square, χ2=9.636, p=0.002) (**Figure 3**). No associations were observed between the number of iron-positive axons and other axonal injury markers: β-APP, which indicates disrupted fast axonal transport;^27^ SMI-32, which identifies disrupted slow axonal transport and phosphorylation abnormalities;^28^ and NPY-Y1R, which is a marker of Wallerian degeneration.^29,30^ (**Figure 3**, **Supplementary Data 6**).

### Iron-positive microglia-macrophage in the non-lesional white matter

Microglia-macrophages commonly demonstrated intense nuclear and cytoplasmic Turnbull staining, often in close proximity to vessels. Iron-positive microglia-macrophages frequently displayed dysmorphic morphological features (**Figure 1**).^12^ In highly inflammatory cases, sparse foamy macrophages were observed near vessels in 23.8% (30/126) samples (**Figure 1E**).

A striking increase in the proportion of iron-positive glial cells with microglia-macrophage morphology was observed in MS (0.27, CI95% 0.18 – 0.36) compared to controls (0.007, CI95% −0.006 – 0.021; Wald χ2=30.63, p<0.001). In the same cohort, Iba-1 was only moderately increased in MS (630, CI95% 510 – 751 10^3^ positive pixels/µm2) compared to controls (373.5, CI95% 257 – 490 10^3^ positive pixels/µm2; Wald χ2=9.07, p=0.003) and correlated with the iron-positive microglia-macrophage score (rho 0.376, p<0.001). A 6-fold increase in iron-positive microglia-macrophages was confirmed when correcting for Iba-1 (0.27, CI95% 0.17 – 0.36 vs 0.077, CI95% 0.004 – 0.15; Wald χ2=9.76, p=0.002). This finding was replicated by double labelling of iron with Iba-1 in a subgroup of 15 cases, in which a near seven-fold increase in the average number of Iba-1 & Turnbull double-positive cells was found between MS and controls, as well as in the relative proportion of Iba-1 & Turnbull double-positive microglia-macrophage on total Iba-1-positive cells was observed (Wald χ2=6.009, p=0.014 and Wald χ2=5.800, p=0.016; respectively) (**Figure 4**, **Supplementary Data 7**).

**Figure 4:**
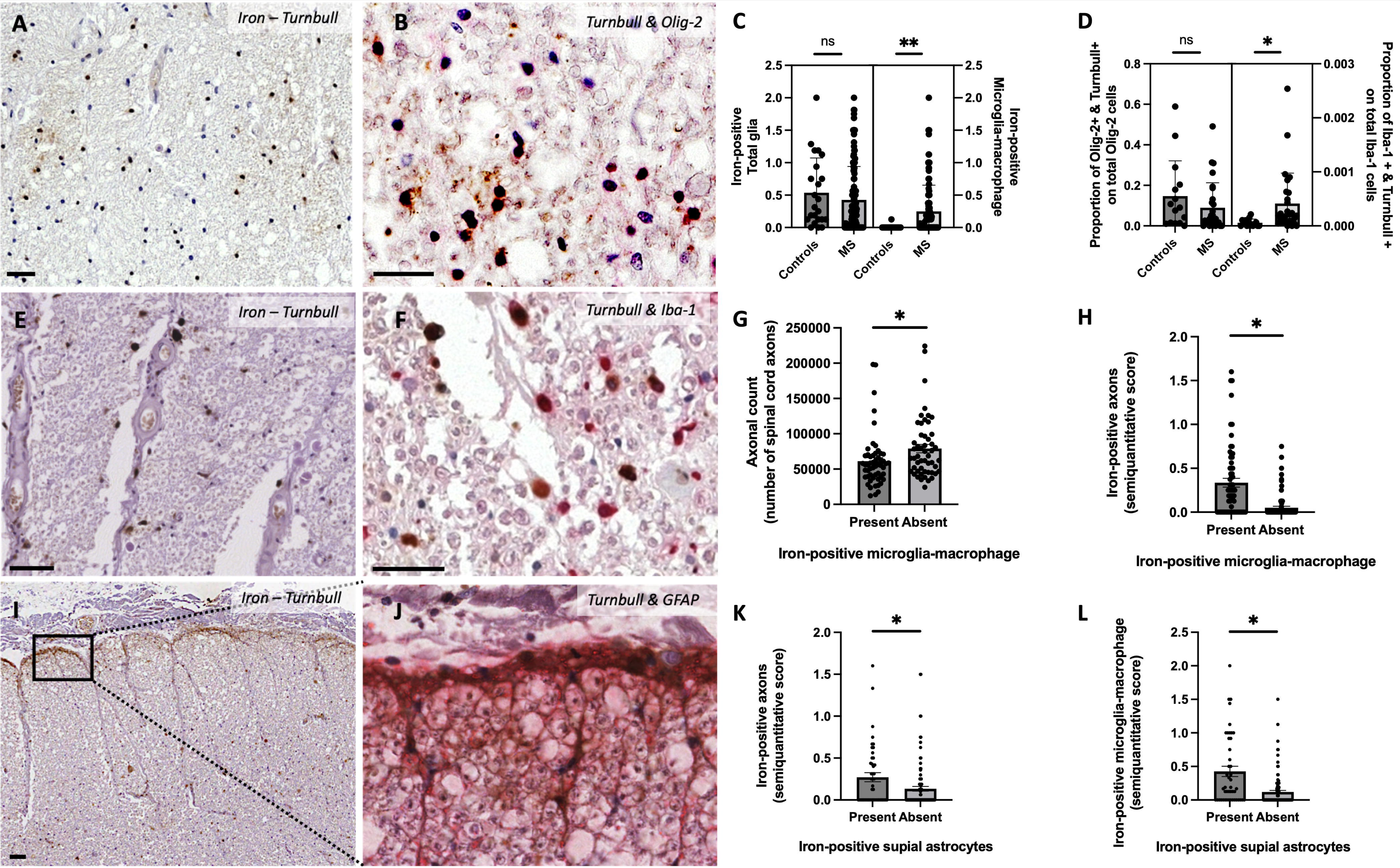
Iron accumulates in non-lesional WM microglia-macrophages and subpial astrocytes in MS. (A-D) Iron-rich oligodendrocytes are morphologically characterized by small and dense nuclear iron surrounded by thin iron-positive processes (A), and express Olig-2 (B). The density of iron-positive glial cells, which predominantly showed oligodendroglial morphology, did not differ between MS and controls (C), a finding confirmed in a subset of 10 MS and 5 control cases by counting the number of co-labelled Turnbull and Olig-2 glial cells (D). Iron-positive activated microglia-macrophages, characterized by large ameboid or dysmorphic morphology (E) and Iba-1 expression (F), were increased in MS compared to control (C, morphological semiquantitative score) as well as the proportion of iron-positive microglia-macrophages on total microglia-macrophages (D, Turnbull and Iba-1 double-labelling). The presence of iron-positive microglia-macrophages in the non-lesional WM was associated with lower axonal counts (G) and increased iron accumulation in the axons (H). In the subpial area, a thick fibrillar pattern of iron staining was observed (I), particularly in MS cases, and showed double labelling with the astrocyte marker GFAP (J). MS cases with subpial astrocytic iron showed higher iron accumulation also in microglia-macrophages (K) and axons (L). Scale bar corresponds to 20µm.

Iron-positive glial cells with microglia-macrophage morphology correlated with higher iron-positive axons (Wald χ2=12.86, p<0.001) and tended to correlate with lower axonal counts (Wald χ2=3.24, p=0.072), after correction for cord level.

Interestingly, similar to iron-positive axons, iron-positive microglia-macrophage showed higher scores in the posterior aspect of the lateralal funiculi in both MS and controls (Friedman’s p<0.001). No effect of level was observed in either MS or controls.

### Iron-positive subpial astrocytes

Subpial astrocytes were more commonly found in MS cases compared to controls, often presenting with hypertrophic features and creating a dense fibrillar network which delineates a high proportion of the spinal cord perimeter.

A subpial band of iron-positive astrocytes was detected in 34.4% of MS cases and in 4.8% of controls (χ2 7.51, p=0.006). The extent of the subpial rim was over 6-fold higher in MS compared with controls, with an average score of 0.46 (CI95% 0.3 – 0.61) vs. 0.075 (CI95% −0.06 – 0.21; Wald χ2 13.3, p<0.001).

Higher iron-positive subpial astrocytes were associated with higher iron-positive axons (Wald χ2 7.85, p=0.005), glia (Wald χ2 4.3, p=0.038), and microglia-macrophages (Wald χ2 5.29, p=0.021), after correction for cord level. No association between iron-positive astrocyte and cord area or axonal counts was detected. Iron-positive subpial astrocyte count did not differ between cord levels. In close proximity to subpial hypertrophic astrocytes, few small iron-positive axons were commonly found between the thick astrocyte processes (**Figure 1H**).

### Iron-positive oligodendrocytes in non-lesional white and grey matter

No significant difference was observed in the density of total iron-positive glial cells that mostly resembled oligodendrocytes in the non-lesional WM between MS (0.41, CI95% 0.29 – 0.53) and controls (0.54, CI95% 0.27 – 0.82; p>0.3). Using double labelling of Turnbull with Olig-2 in a cohort of 10 MS and 5 control cases, the total number of Olig-2-positive cells, iron-positive oligodendrocytes, or the proportion of iron-positive oligodendrocyte on total Olig-2 positive cells did not differ between groups (**Figure 4**, Supplementary data 4).

In GM, 76.2% of controls displayed a large cluster of iron-positive oligodendrocytes in the dorsal root entry zone and dorsal horn compared to 57.3% of the MS cases. Moreover, 52.4% of controls showed iron-positive oligodendrocytes across the entire GM area compared to 24.4% of MS cases. Overall, a reduced prevalence of iron-positive oligodendrocytes was observed in the GM in MS (1.26, CI95% 0.86 – 1.66 vs. 0.8, CI95% 0.6 – 0.99; Wald χ2 4.13, p=0.042).

### 3.3 Iron in spinal cord demyelinating lesions

Sixty-seven WM demyelinating lesions were identified in our spinal cord cohort, of which 13.4% (9/67) were active, 61.2% (41/67) were mixed active-inactive, and 25.4% (17/67) inactive. None of these WM lesions showed an iron-positive rim of microglia-macrophages at the lesion edge (Figure 5A-D). Of note, characterisation of mixed active-inactive lesions was extended to include an additional 10 mixed active-inactive lesions from a larger cohort of 119 spinal cord samples,^31^ with no rim being identified in this additional cohort.

**Figure 5:**
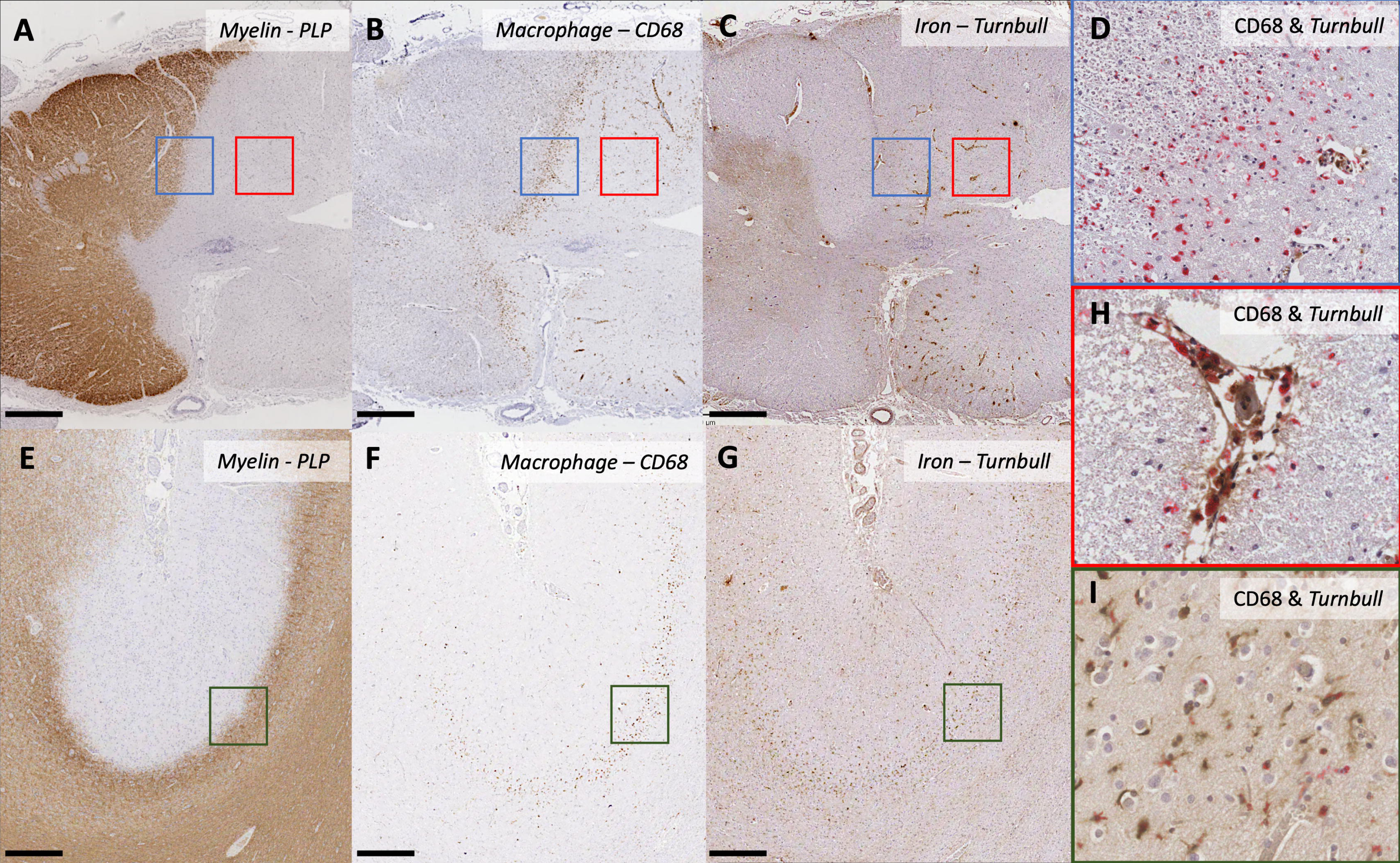
Iron accumulates in perivascular macrophages in both acute and chronic active lesions. **(A)** Myelin (PLP) staining reveals a demyelinating plaque in a cervical spinal cord section from a progressive case of multiple sclerosis (MS). **(B)** Inflammation characterized by microglia-macrophage (CD68+) infiltration exhibits a typical rim of intense inflammatory activity at the border of the plaque. **(C)** Minimal accumulation of non-heme iron, as indicated by DAB-enhanced Turnbull staining, is observed in the microglia-macrophages at the rim of the plaque. (Magnification in **D** for double labelling with Turnbull/DAB in brown and CD68/Vector Red in red). **H:** Conversely, an intense iron accumulation is observed in perivascular microglia-macrophages within the plaque’s edge. (Magnification in **H** for double labelling with Turnbull/DAB in brown and CD68/Vector Red in red). In the mesiofrontal motor cortex samples from the same autopsy cohort show iron-rimmed demyelinating lesions. **(E)** Myelin (PLP) staining shows a type III cortical lesion. **(F)** A rim of microglia-macrophage (CD68+) is seen at the plaque border. **(G)** A reduction in iron intensity is seen in the lesion core, with a rim of iron-positive microglia-macrophages at the plaque border. (Magnification in **I** for double labeling with Turnbull/DAB in brown and CD68/Vector Red in red). The scale bar corresponds to 800µm.

While parenchymal macrophages, forming the rim at the edge of the lesion, did not show iron-positivity, vessel-associated macrophages showed iron positivity in areas of active inflammation (Figure 5E). Perivascular foamy macrophages with a granular Turnbull-positive cytoplasmic stain formed rich perivascular cuffs in areas of active demyelination in both acute and mixed active lesions (Figure 1D). These perivascular cells were observed in 55.6% (5/9) of acute, 51.2% (21/41) of mixed active-inactive, and 5.9% (1/17) of inactive lesions.

Occasionally, iron-positive axons with swollen morphology were observed in active demyelinating lesions, as previously described.^32^ In particular, iron-positive axons were identified in two active lesions and at the active border of 7 mixed active-inactive lesions.

Few hypertrophic astrocytes with thick iron-rich processes were observed in the gliotic core of inactive or mixed active-inactive lesions. In one case, iron-positive axons were found in close proximity to iron-positive astrocytes in the plaque (**Figure 1I**).

### 3.4 Iron distribution in the non-lesional mesiofrontal motor cortex

Thirty-eight mesiofrontal brain sections from 36 MS cases and 3 controls were included in the study. The primary motor cortex showed prominent and patchy Turnbull staining of myelin fibre compared to the spinal cord, consistent with the higher iron concentration reported in this brain region.^25^

Subcortical WM exhibited a tigroid Turnbull staining pattern associated with vascular structures (**Figure 1J**), with clusters of iron-positive oligodendrocytes found along vessels (**Figure 1K**). Iron-positive microglia-macrophages were occasionally seen, particularly within or in close proximity to vessels.

In the cortical GM, deeper layers (IIIb-VI) showed consistently high Turnbull reactivity while layer II typically had a low iron content. Most iron-positivity was associated with cells with oligodendroglial morphology and some perivascular microglia-macrophages. Bundles of iron-stained myelin projecting from layers III-V into subcortical WM was a consistent feature. No definite axonal positivity for Turnbull was identified in our motor cortical cohort. Interestingly, layer I showed variable Turnbull staining between cases, characterised by both iron-positive oligodendrocytes and subpial fibrillar staining.

### 3.4 Iron in grey- and subcortical white-matter demyelinating lesions in the mesiofrontal cortex

In the MS motor cortex, we identified 43 lesions, specifically six type I lesions, five type II lesions, 21 type III lesions, and 11 type IV lesions. Five mixed active-inactive lesions were identified in subcortical WM.

Turnbull staining was reduced at the lesion core. A rim of iron-positive microglia-macrophage was observed at the border of cortical lesions, a finding more frequently observed in deeper cortical layers, especially where subcortical WM was involved. Specifically, iron rimmed lesions were observed in 20.9% of all cortical lesions studied, being present in 50% of type I lesions, 19.05% of type III lesions, 18.2% of type IV lesions, and in 80% of subcortical lesions (**Figure 5**).

### 3.6 ICP-OES quantification of non-haem iron

Quantification of total non-haem iron was performed in the spinal cord cohort. A sodium citrate pH 1.0 buffer was used to extract non-haem iron from the tissue sections, according to an optimised method.^19^ Iron concentration in the citrate buffer was quantified by ICP-OES. The efficacy of the buffer in extracting iron was evidenced by the ability of the buffer to completely abolish the Turnbull signal (**Figure 6**, **Supplementary Data 1**).

**Figure 6:**
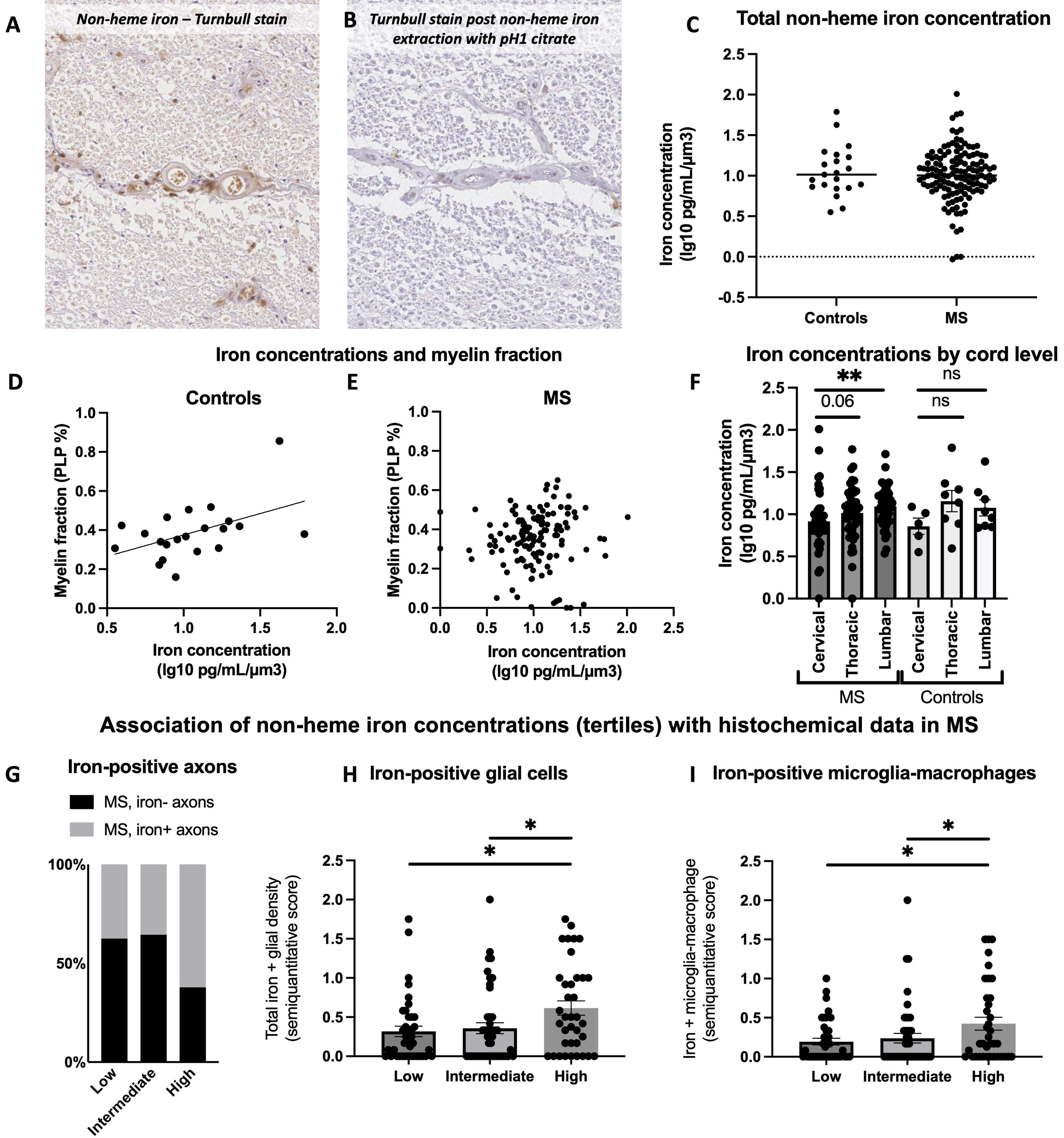
Non-heme iron concentrations relate to myelin fraction in controls, while showing a disto-proximal gradient and an association with iron-positive axons and microglia-macrophages in MS. Non-heme iron has been extracted from spinal cord sections using a sodium citrate pH 1.0 solution. DAB-enhanced Turnbull staining of two adjacent sections which have previously incubated for 72h in water (A) or acid citrate (B) shows the ability of acid citrate to extract non-heme iron. No difference in total iron concentration was observed between MS and controls (C). Iron concentration was strictly related to myelin fraction in controls (PLP area coverage; rho 0.48, p=0.028) but not in MS cases (p>0.3). On the contrary, iron concentrations increased distally in the cord in MS cases (Kruskal-Wallis p=0.027, p=0.013) but not in controls (p>0.2). When dividing iron concentrations in tertiles, MS samples with higher iron content had a higher prevalence of iron-positive axons (Chi-square 6.93 p =0.031), and showed higher total iron-positive glia (Kruskal-Wallis p=0.027) and iron-positive microglia-macrophages (Kruskal-Wallis p=0.03) compared to the intermediate and low iron content samples.

No difference in total non-haem iron concentration was observed between MS and controls. In controls, iron concentration correlated with higher myelin fraction (Wald χ2 26.45, p<0.001) while no such association was found between iron concentrations and myelin fraction in MS, once correcting for cord level. On the contrary, iron concentration was greater in the distal compared to proximal levels of the spinal cord in MS (Wald χ2 4.4, p=0.036) with no such difference found in controls, once correcting for myelin fraction (**Figure 6**). These findings apparently mirror the disto-proximal gradient of axonal staining observed in MS.

When binning iron concentration into tertiles (high, intermediate, low iron concentration), higher iron concentration associated with presence of iron-positive (A) axons (Wald χ2 3.87, p=0.049), (B) total iron-positive glia (Wald X2 4.85, p=0.028) and (C) microglia-macrophage scores (Wald χ2 4.26, p=0.039) in MS. No association was found with subpial astrocytes. Importantly, high iron concentration also associated with reduced axonal counts (Wald χ2 12.73 p<0.001) in MS. No association was found with cord area. All analyses were corrected for cord level (**Figure 6**).

## 4. Discussion

We present a comprehensive and systematic characterisation of iron distribution in the MS spinal cord and illuminate several important findings. First, iron-rimmed lesions are not detected in the MS spinal cord compared to the motor cortex where typical iron-rims are found at the edge of cortical and subcortical lesions. Given that iron content is notably low in the spinal cord compared to the motor cortex/subcortical area, our findings suggest that PRLs may be a biomarker of lesion activity dependent on iron content of local myelin,^25^ rather than being a specific marker of severe pathology. Second, iron is diffusely and aberrantly deposited in non-lesional spinal cord tissue with pronounced accumulation in axons, dysmorphic microglia-macrophages, and subpial astrocytes. Third, iron accumulation in axons shows a predilection for the motor tract with a disto-proximal gradient of involvement being most pronounced in the lumbar spinal cord; this anatomical distribution is unique to pathological studies and matches the predominant lower limb motor disturbance often encountered in progressive MS. Finally, total iron concentration and iron-positive axons associate with markers of neurodegeneration (axonal counts and cord atrophy), adding further credence to a role for iron in disability accumulation in MS. The conflation of our findings points to aberrant iron deposition in the MS spinal cord as a key feature of neurodegeneration, and by extension, disability accumulation, in progressive MS.

### 4.1 Aberrant iron distribution in the MS spinal cord

In health, most CNS iron is stored in myelin and oligodendrocytes where it is required for myelin synthesis and energy production.^33^ As systematically assessed by Morris and colleagues in 1992, iron levels vary between CNS regions, being highest in deep GM nuclei and subcortical WM, and lowest in the spinal cord.^25^ Within the spinal cord, most iron is contained in oligodendrocytes of laminae I-III and the fasciculus of Lissauer.^25^ Our findings in control motor cortical and spinal cord sections match these descriptions. We expand these findings by demonstrating that total non-haem iron content, as measured by ICP-OES, strictly correlates with myelin fraction in controls.

In MS, total iron concentration does not differ from controls but is independent of myelin fraction, suggesting a shift of iron from myelin/oligodendrocytes to other CNS compartments. The striking iron reactivity observed in MS spinal cord axons, activated microglia-macrophages, and subpial astrocytes, all associating with higher non-haem iron content in MS, not only supports such a redistribution but also validates our semiquantitative scoring method of the Turnbull stain. What is more, the fact that total non-haem iron concentration relates to axonal loss, a known correlate of disability in progressive MS. Overall, these data demonstrate an aberrant distribution of iron in the MS spinal cord that has relevance to disability accumulation in progressive disease.

The notion of aberrant iron distribution in MS has already been suggested by prior pathological and imaging studies. Iron content has been reported to be reduced in perilesional WM in cases who are older^34^ and with longer disease duration.^12^ In contrast, iron content has been shown to be slightly increased in non-lesional deep GM, independent of age, and associates with greater oxidative stress and neuro-axonal loss when present in deep lesional GM.^13^ Leveraging the paramagnetic properties of iron, MRI studies have demonstrated approximately a 30% increase in iron concentration in deep GM nuclei in MS compared to controls,^35^particularly in cases with longer disease duration^36^ and greater disability, regardless of atrophy.^37^ Interestingly, these clinically relevant radiographic changes do not simply reflect an increase in total iron content, which may in fact be reduced,^38^ at least in certain regions.^37^ Instead, the imaging findings support a shift in the cellular distribution of iron in MS,^38^ in line with our spinal cord findings. Moreover, the topographical specificity of iron changes in MS highlights the complexity of iron metabolism in the disease and underscores the need for a systematic assessment of iron along the neuraxis. Our findings point to heretofore undescribed changes in iron metabolism in the MS spinal cord, which is arguably the most relevant area for disability progression.^39^

### 4.2 Iron accumulation in spinal cord axons relates to neurodegeneration and oxidative stress in MS

Here we describe selective iron accumulation in spinal cord axons in MS, a pattern that associates with smaller cord area and axonal loss, both known determinants of disability in progressive MS.^39^ The preferential accumulation of iron in the motor pathway in a disto-proximal gradient intriguingly aligns with the predominant disturbance of motor function in the lower limbs characteristic of progressive MS, which is unlike any other known pathological signatures described to date.

Iron accumulation within axons has previously been observed in the context of axonal swellings in active lesions – a finding we also report in this autopsy cohort.^32^ We extend these findings by showing widespread and symmetric iron reactivity within notional LCST tracts in non-lesional WM outside of areas of acute axonal injury or Wallerian degeneration. Prior studies may not have captured these observations due to restricted tissue sampling (i.e. brain only) and technical factors, such as time in formalin, which is known to extract iron.^40,41^

Ample clinical, radiographic and pathological data suggest that neurodegeneration in the spinal cord occurs independently of demyelination,^3,39,42,43^ with the underlying mechanisms being poorly understood. Oxidative stress and mitochondrial dysfunction contribute to neuro-axonal energy failure and have been considered important contributors of disease progression, with iron accumulation possibly catalysing this process.^7,8^ Upper motor neurons, whose long projection axons extend over a metre from the motor cortex down the length of the spinal cord, are particularly vulnerable in a multifocal disease like MS; and in fact, these nerve fibres have been shown to contain mitochondria deficient in respiratory chain enzymes and mitochondrial DNA deletions suggestive of accelerated biological aging.^8,44,45^ A recent landmark study demonstrated that cells exposed to different types of senescence-inducing insults accumulate iron, both in ferritin-bound and labile forms, which predispose to oxidative stress and to acquiring a senescence-associated secretory profile (SASP) that propagates senescence in surrounding cells.^46^ The observed disto-proximal gradient of iron accumulation in the spinal cord axons, confirmed by ICP-OES quantification, is highly suggestive of an energetic failure or metabolic dysfunction pathogenesis. It has also been shown that spinal cord mitochondria are more vulnerable to calcium overload than brain mitochondria,^47^ which may support an inherent vulnerability to energy failure and oxidative stress in this clinically eloquent CNS region. Iron-positive axons associated with axonal expression of oxidised phospholipids which are labelled by E06, an antibody that targets oxidised phospholipids and has been reported as a marker of oxidative stress, and of iron-related programmed cell death aka ferroptosis.^48,49^ The lack of association with β-APP and SMI-32 further supports that iron-axons are in a state of chronic oxidative stress - rather than acute axonal injury.^50^ The preferential accumulation of iron in the anterior and lateral corticospinal tracts axons and the symmetrical involvement of two hemicords of non-lesional white matter suggests a specific vulnerability of motor axons for iron overload. The length and elevated energy demand of these axons may be a plausible predisposing factor to mechanisms of energy failure and senescence. It is also possible to speculate that a higher physiological concentration of iron in motor axons may contribute to the ability to more sensitively detect iron overload in these tracts due to a “floor effect” or threshold of the histochemical method used.

### 4.3 Diffuse infiltration of iron-loaded dysmorphic microglia-macrophages in the non-lesional MS spinal cord

We observed a selective and marked increase in the number of iron-loaded microglia-macrophages across the non-lesional WM, in excess of what would be expected based on total microglia-macrophage infiltration. These cells often displayed dysmorphic morphology and were associated with iron-positive axons.

Microglia-macrophage infiltration in non-lesional WM is a key characteristic of progressive MS pathology, which has been shown to relate to neurodegeneration and disability, as evidenced by both pathological and imaging studies.^8,51–53^ Altered iron metabolism is a known feature of infiltrating microglia-macrophages in progressive MS,^12,17^ even in non-lesional tissue.^13,53–55^ Senescent microglia-macrophages accumulate iron in normal ageing and other neurodegenerative conditions,^56–58^ develop a degenerative/dysmorphic morphology, promote the expression of ferroptosis-related genes, contribute to mitochondrial damage, oxidative stress, and a self-propagating pro-inflammatory secretory phenotype.^55–59^ Our finding that iron-loaded microglia-macrophages closely associate with iron accumulation in axons suggests a complex interplay between microglia and their surrounding microenvironment; this is supported by the observations that iron-loaded microglia can facilitate ferroptosis in surrounding neuronal cells,^59^ and that iron overload can propagate and induce senescence in adjacent cells.^46^ Overall, our work supports iron dysmetabolism of microglia-macrophages and its impact on the local microenvironment, as being a key feature of spinal cord pathology in progressive MS.

What is the provenance of iron in both ageing and MS? Multiple sources are implicated, including blood–brain barrier permeability, inflammation, and myelin phagocytosis.^17,56,57^ Notably, we found that iron-loaded microglia-macrophages were more prevalent in the lateral columns—where the longest perforating arterioles run—and tended to accumulate in perivascular areas. Moreover, the fact that increased iron within the microglia-macrophage compartment did not associate with reduced iron-positive oligodendrocytes or myelin fraction, also supports a possible vascular origin.^17^ Recent *in vitro* data have demonstrated that vascular leakage and subsequent iron overload may induce senescence in macrophages, leading to further iron accumulation and promotion of similar changes in surrounding cells, creating a self-propagating cycle.^46^ It is plausible to speculate that such mechanisms also occur in the MS brain, where early blood-brain barrier damage contributes to iron dysmetabolism and overload that accelerates immune and neural cell senescence, which, in turn, drives relentless neurodegeneration that continues even after the initial triggering event has subsided. Could this explain why even potent immunomodulatory/suppressive and neuroprotective therapies in MS are ineffective in treatment the progressive phase of the disease?

### 4.4 Iron accumulates in hypertrophic subpial astrocytes in MS

We found that hypertrophic, iron-rich subpial astrocytes were frequently present in MS. Astrocytes have strong iron transport capacity and are relatively resistant to oxidative stress.^60^ Iron-positive astrocytes have been previously described in the core of chronic MS lesion – a finding repliced in our study - and were considered to have protective effects in buffering iron excess.^12,34^ We show that subpial iron astrocytes correlate with iron accumulation in axonal and microglia/macrophage compartments, suggesting their presence identifies cases with highest iron overload. Interestingly, the high intensity of iron in these cells make them amenable to *in vivo* MRI tracking as suggested by a recent 7T imaging study on people with MS, which could serve as a biomarker of the iron findings presented herein.^11^

### 4.5 Iron-rimmed lesions are specific to brain MS pathology and are absent in the spinal cord

Iron-rims were frequently found in both cortical and subcortical WM lesions, yet were notably absent in the MS spinal cord. Our study also provides data on the prevalence of iron-rimmed lesions in the cortex, which has not been systematically reported to date.

Iron-rimmed lesions in the brain are known to strongly correlate with disability outcomes in MS.^10,61^ However, no prior pathological study has examined iron in spinal cord lesions, and its detection using MRI has long been thought to be limited by technical factors. In our large cohort of MS spinal cord samples, remarkably, no iron-rimmed lesions were identified. This contrasts with the fact that iron-rimmed lesions were detected in mesiofrontal cortical brain samples from the same cases. The only study describing paramagnetic signals in the MS spinal cord has shown that the most intense paramagnetic signal is in the subpial area and at the grey-white matter boundaries—a finding that strikingly aligns with our qualitative observations, particularly regarding subpial astrocytes.^11^ The topographical variability in myelin iron-content between CNS regions may also explain the higher prevalence of iron-rimmed cortical lesions we observed in the mesio-frontal cortex compared to prior studies looking at the whole cortical ribbon.^62^

There is ongoing debate over whether the iron accumulation in microglia-macrophages at the lesion edge originates from myelin debris or the vasculature. The latter hypothesis is supported by the expression of haemoglobin-haptoglobin receptor (CD163) and heme-oxygenase-1 protein in microglial cells at edge of lesions.^17^ Comparing lesions from CNS regions with varying iron concentrations, such as the spinal cord and subcortical WM, uniquely places us to disentangle these two iron sources. We show that parenchymal microglial infiltration at the lesion edge decreases in iron content when transitioning from regions with high myelin iron (i.e. subcortical and type I cortical lesions) to regions with low myelin iron (i.e. spinal cord). However, perivascular and intravascular macrophages in highly inflammatory areas of the plaque consistently demonstrated iron-reactivity in both cortical and spinal cord lesions, supporting the role for vascular iron influx.

## 5. Strengths and limitations

Several *in vitro*, genetic, imaging, and CSF-proteomic data have demonstrated a role for iron dysmetabolism in MS. However, a systematic description of iron distribution in non-lesional MS tissue had not been conducted to date and little was known about iron in the spinal cord. We had access to a large cohort of post-mortem spinal cord and brain samples that enable us to address this gap and reveal striking patterns of iron distribution relevant to neurodegeneration heretofore not described.

There are inherent limitations to the study of post-mortem material. First, pathology provides a static view of dynamic phenomena and does not allow determination of mechanisms responsible for iron accumulation, its upstream contributors or downstream consequences (e.g. axonal damage, oxidative stress, senescence). I*n vitro* studies have shown that iron dysmetabolism and senescence are intertwined mechanisms of a same self-propelling cycle,^46,56^ which require further study in MS. Second, iron characterisation was performed using histochemical methods not suitable for an exact quantification of iron content. We addressed this limitation, in part, by validating our findings through a novel ICP-OES quantification of non-haem iron in matched samples. Third, the process of formalin fixation is known to reduce iron levels,^41^ which may have reduced our sensitivity to detect iron in both controls and MS.

In summary, our findings demonstrate widespread, tract-specific, and length-dependent aberrant iron distribution in the MS spinal cord that relates to neurodegeneration independent of demyelination. The preferential accumulation of iron in motor tract axons and its association with oxidative stress, axonal loss, and spinal cord atrophy, suggests a role for iron redistribution in driving disease progression in MS. Moreover, the absence of iron-rimmed lesions in the spinal cord contrasts with their presence in cortical and subcortical areas highlighting regional specificity; these findings when combined with the striking relationship between non-lesional iron and neurodegenerative outcomes in the MS spinal cord suggest that imaging markers of iron beyond PRLs are urgently needed. Future work is needed to understand how iron dysmetabolism and cellular senescence in the MS spinal cord contribute to disease progression, with the goal of identifying pathways amenable to therapeutic intervention.

## Aknowledgements

We acknowledge the Laboratório de Análises/REQUIMTE/LAQV for the acquisition of the ICP-OES data

## Bibliography

1. Dow RS. VASCULAR PATTERN OF LESIONS OF MULTIPLE SCLEROSIS. Arch Neurol Psychiatry. 1942;47(1):1. doi:10.1001/archneurpsyc.1942.02290010011001

2. Filippi M, Bar-Or A, Piehl F, et al. Multiple sclerosis. Nat Rev Dis Primer. 2018;4(1):1–27. doi:10.1038/s41572-018-0041-4

3. Confavreux C, Vukusic S. Natural history of multiple sclerosis: a unifying concept. Brain J Neurol. 2006;129(Pt 3):606–616. doi:10.1093/brain/awl007

4. Confavreux C, Vukusic S, Moreau T, Adeleine P. Relapses and progression of disability in multiple sclerosis. N Engl J Med. 2000;343(20):1430–1438. doi:10.1056/NEJM200011163432001

5. Leray E, Yaouanq J, Le Page E, et al. Evidence for a two-stage disability progression in multiple sclerosis. Brain. 2010;133(7):1900–1913. doi:10.1093/brain/awq076

6. Amato MP, Fonderico M, Portaccio E, et al. Disease-modifying drugs can reduce disability progression in relapsing multiple sclerosis. Brain. 2020;143(10):3013–3024. doi:10.1093/brain/awaa251

7. Mahad DH, Trapp BD, Lassmann H. Pathological mechanisms in progressive multiple sclerosis. Lancet Neurol. 2015;14(2):183–193. doi:10.1016/S1474-4422(14)70256-X

8. Kuhlmann T, Moccia M, Coetzee T, et al. Multiple sclerosis progression: time for a new mechanism-driven framework. Lancet Neurol. 2023;22(1):78–88. doi:10.1016/S1474-4422(22)00289-7

9. Bagnato F, Hametner S, Yao B, et al. Tracking iron in multiple sclerosis: a combined imaging and histopathological study at 7 Tesla. Brain. 2011;134(12):3602–3615. doi:10.1093/brain/awr278

10. Dal-Bianco A, Grabner G, Kronnerwetter C, et al. Slow expansion of multiple sclerosis iron rim lesions: pathology and 7 T magnetic resonance imaging. Acta Neuropathol (Berl). 2017;133(1):25–42. doi:10.1007/s00401-016-1636-z

11. Clarke MA, Witt AA, Robison RK, et al. Cervical spinal cord susceptibility-weighted MRI at 7T: Application to multiple sclerosis. NeuroImage. 2023;284:120460. doi:10.1016/j.neuroimage.2023.120460

12. Hametner S, Wimmer I, Haider L, Pfeifenbring S, Brück W, Lassmann H. Iron and neurodegeneration in the multiple sclerosis brain. Ann Neurol. 2013;74(6):848–861. doi:10.1002/ana.23974

13. Haider L, Simeonidou C, Steinberger G, et al. Multiple sclerosis deep grey matter: the relation between demyelination, neurodegeneration, inflammation and iron. J Neurol Neurosurg Psychiatry. 2014;85(12):1386–1395. doi:10.1136/jnnp-2014-307712

14. Stephenson E, Nathoo N, Mahjoub Y, Dunn JF, Yong VW. Iron in multiple sclerosis: roles in neurodegeneration and repair. Nat Rev Neurol. 2014;10(8):459–468. doi:10.1038/nrneurol.2014.118

15. Giordano A, Stridh P, Preziosa P, et al. Genetic variation in HIF1A is associated with smoldering inflammation and disease progression in Multiple Sclerosis. Published online March 16, 2024:2024.03.15.24304290. doi:10.1101/2024.03.15.24304290

16. Magliozzi R, Hametner S, Facchiano F, et al. Iron homeostasis, complement, and coagulation cascade as CSF signature of cortical lesions in early multiple sclerosis. Ann Clin Transl Neurol. 2019;6(11):2150–2163. doi:10.1002/acn3.50893

17. Hofmann A, Krajnc N, Dal-Bianco A, et al. Myeloid cell iron uptake pathways and paramagnetic rim formation in multiple sclerosis. Acta Neuropathol (Berl). 2023;146(5):707–724. doi:10.1007/s00401-023-02627-4

18. Meguro R, Asano Y, Odagiri S, Li C, Iwatsuki H, Shoumura K. Nonheme-iron histochemistry for light and electron microscopy: a historical, theoretical and technical review. Arch Histol Cytol. 2007;70(1):1–19. doi:10.1679/aohc.70.1

19. Van Duijn S, Nabuurs RJA, Van Duinen SG, Natté R. Comparison of Histological Techniques to Visualize Iron in Paraffin-embedded Brain Tissue of Patients with Alzheimer’s Disease. J Histochem Cytochem. 2013;61(11):785–792. doi:10.1369/0022155413501325

20. DeLuca GC, Ebers GC, Esiri MM. Axonal loss in multiple sclerosis: a pathological survey of the corticospinal and sensory tracts. Brain J Neurol. 2004;127(Pt 5):1009–1018. doi:10.1093/brain/awh118

21. DeLuca GC, Alterman R, Martin JL, et al. Casting light on multiple sclerosis heterogeneity: the role of HLA-DRB1 on spinal cord pathology. Brain J Neurol. 2013;136(Pt 4):1025–1034. doi:10.1093/brain/awt031

22. Bankhead P, Loughrey MB, Fernández JA, et al. QuPath: Open source software for digital pathology image analysis. Sci Rep. 2017;7(1):16878. doi:10.1038/s41598-017-17204-5

23. Kuhlmann T, Ludwin S, Prat A, Antel J, Brück W, Lassmann H. An updated histological classification system for multiple sclerosis lesions. Acta Neuropathol (Berl). 2017;133(1):13–24. doi:10.1007/s00401-016-1653-y

24. Bø L, Vedeler CA, Nyland HI, Trapp BD, Mørk SJ. Subpial demyelination in the cerebral cortex of multiple sclerosis patients. J Neuropathol Exp Neurol. 2003;62(7):723–732. doi:10.1093/jnen/62.7.723

25. Morris CM, Candy JM, Oakley AE, Bloxham CA, Edwardson JA. Histochemical Distribution of Non-Haem Iron in the Human Brain. Cells Tissues Organs. 1992;144(3):235–257. doi:10.1159/000147312

26. Spatz H. Über den eisennachweis im gehirn, besonders in zentren des extrapyramidal-motorischen systems. I. Teil. Z Für Gesamte Neurol Psychiatr. 1922;77(1):261–390. doi:10.1007/BF02865844

27. Ferguson B. Axonal damage in acute multiple sclerosis lesions. Brain. 1997;120(3):393–399. doi:10.1093/brain/120.3.393

28. Schirmer L, Antel JP, Brück W, Stadelmann C. Axonal Loss and Neurofilament Phosphorylation Changes Accompany Lesion Development and Clinical Progression in Multiple Sclerosis. Brain Pathol. 2011;21(4):428–440. doi:10.1111/j.1750-3639.2010.00466.x

29. Dziedzic T, Metz I, Dallenga T, et al. Wallerian Degeneration: A Major Component of Early Axonal Pathology in Multiple Sclerosis. Brain Pathol. 2010;20(5):976–985. doi:10.1111/j.1750-3639.2010.00401.x

30. Pesini P, Kopp J, Wong H, Walsh JH, Grant G, Hökfelt T. An immunohistochemical marker for Wallerian degeneration of fibers in the central and peripheral nervous system. Brain Res. 1999;828(1-2):41–59. doi:10.1016/s0006-8993(99)01283-4

31. Waldman AD, Catania C, Pisa M, Jenkinson M, Lenardo MJ, DeLuca GC. The prevalence and topography of spinal cord demyelination in multiple sclerosis: a retrospective study. Acta Neuropathol (Berl). 2024;147(1):51. doi:10.1007/s00401-024-02700-6

32. Craelius W, Migdal MW, Luessenhop CP, Sugar A, Mihalakis I. Iron deposits surrounding multiple sclerosis plaques. Arch Pathol Lab Med. 1982;106(8):397–399.

33. Connor JR, Menzies SL. Relationship of iron to oligondendrocytes and myelination. Glia. 1996;17(2):83–93. doi:10.1002/(SICI)1098-1136(199606)17:2<83::AID-GLIA1>3.0.CO;2-7

34. Popescu BF, Frischer JM, Webb SM, et al. Pathogenic implications of distinct patterns of iron and zinc in chronic MS lesions. Acta Neuropathol (Berl). 2017;134(1):45–64. doi:10.1007/s00401-017-1696-8

35. Ge Y, Jensen JH, Lu H, et al. Quantitative Assessment of Iron Accumulation in the Deep Gray Matter of Multiple Sclerosis by Magnetic Field Correlation Imaging. Am J Neuroradiol. 2007;28(9):1639–1644. doi:10.3174/ajnr.A0646

36. Elkady AM, Cobzas D, Sun H, Blevins G, Wilman AH. Progressive iron accumulation across multiple sclerosis phenotypes revealed by sparse classification of deep gray matter. J Magn Reson Imaging JMRI. 2017;46(5):1464–1473. doi:10.1002/jmri.25682

37. Zivadinov R, Tavazzi E, Bergsland N, et al. Brain Iron at Quantitative MRI Is Associated with Disability in Multiple Sclerosis. Radiology. 2018;289(2):487–496. doi:10.1148/radiol.2018180136

38. Schweser F, Hagemeier J, Dwyer MG, et al. Decreasing brain iron in multiple sclerosis: The difference between concentration and content in iron MRI. Hum Brain Mapp. 2021;42(5):1463–1474. doi:10.1002/hbm.25306

39. Bischof A, Papinutto N, Keshavan A, et al. Spinal Cord Atrophy Predicts Progressive Disease in Relapsing Multiple Sclerosis. Ann Neurol. 2022;91(2):268–281. doi:10.1002/ana.26281

40. Schrag M, Dickson A, Jiffry A, Kirsch D, Vinters HV, Kirsch W. The effect of formalin fixation on the levels of brain transition metals in archived samples. BioMetals. 2010;23(6):1123–1127. doi:10.1007/s10534-010-9359-4

41. Hametner S, Endmayr V, Deistung A, et al. The influence of brain iron and myelin on magnetic susceptibility and effective transverse relaxation - A biochemical and histological validation study. NeuroImage. 2018;179:117–133. doi:10.1016/j.neuroimage.2018.06.007

42. Weinshenker BG, Bass B, Rice GP, et al. The natural history of multiple sclerosis: a geographically based study. I. Clinical course and disability. Brain J Neurol. 1989;112 ( Pt 1):133–146. doi:10.1093/brain/112.1.133

43. Evangelou N, DeLuca GC, Owens T, Esiri MM. Pathological study of spinal cord atrophy in multiple sclerosis suggests limited role of local lesions. Brain J Neurol. 2005;128(Pt 1):29–34. doi:10.1093/brain/awh323

44. Campbell G, Mahad DJ. Mitochondrial dysfunction and axon degeneration in progressive multiple sclerosis. FEBS Lett. 2018;592(7):1113–1121. doi:10.1002/1873-3468.13013

45. Campbell GR, Ziabreva I, Reeve AK, et al. Mitochondrial DNA deletions and neurodegeneration in multiple sclerosis. Ann Neurol. 2011;69(3):481–492. doi:10.1002/ana.22109

46. Maus M, López-Polo V, Mateo L, et al. Iron accumulation drives fibrosis, senescence and the senescence-associated secretory phenotype. Nat Metab. 2023;5(12):2111–2130. doi:10.1038/s42255-023-00928-2

47. Morota S, Hansson MJ, Ishii N, Kudo Y, Elmér E, Uchino H. Spinal cord mitochondria display lower calcium retention capacity compared with brain mitochondria without inherent differences in sensitivity to cyclophilin D inhibition. J Neurochem. 2007;103(5):2066–2076. doi:10.1111/j.1471-4159.2007.04912.x

48. Van San E, Debruyne AC, Veeckmans G, et al. Ferroptosis contributes to multiple sclerosis and its pharmacological targeting suppresses experimental disease progression. Cell Death Differ. 2023;30(9):2092–2103. doi:10.1038/s41418-023-01195-0

49. Muñoz U, Sebal C, Escudero E, et al. Main Role of Antibodies in Demyelination and Axonal Damage in Multiple Sclerosis. Cell Mol Neurobiol. 2022;42(6):1809–1827. doi:10.1007/s10571-021-01059-6

50. Haider L, Fischer MT, Frischer JM, et al. Oxidative damage in multiple sclerosis lesions. Brain J Neurol. 2011;134(Pt 7):1914–1924. doi:10.1093/brain/awr128

51. Sucksdorff M, Matilainen M, Tuisku J, et al. Brain TSPO-PET predicts later disease progression independent of relapses in multiple sclerosis. Brain. 2020;143(11):3318–3330. doi:10.1093/brain/awaa275

52. Pisa M, Pansieri J, Yee S, et al. Anterior optic pathway pathology in CNS demyelinating diseases. Brain J Neurol. 2022;145(12):4308–4319. doi:10.1093/brain/awac030

53. Zrzavy T, Hametner S, Wimmer I, Butovsky O, Weiner HL, Lassmann H. Loss of “homeostatic” microglia and patterns of their activation in active multiple sclerosis. Brain J Neurol. 2017;140(7):1900–1913. doi:10.1093/brain/awx113

54. van der Poel M, Ulas T, Mizee MR, et al. Transcriptional profiling of human microglia reveals grey–white matter heterogeneity and multiple sclerosis-associated changes. Nat Commun. 2019;10(1):1139. doi:10.1038/s41467-019-08976-7

55. Proto JD, Zhang M, Ryan S, et al. Disrupted microglial iron homeostasis in progressive multiple sclerosis. Published online July 10, 2021:2021.05.09.443127. doi:10.1101/2021.05.09.443127

56. Ward RJ, Zucca FA, Duyn JH, Crichton RR, Zecca L. The role of iron in brain ageing and neurodegenerative disorders. Lancet Neurol. 2014;13(10):1045–1060. doi:10.1016/S1474-4422(14)70117-6

57. Adeniyi PA, Gong X, MacGregor E, et al. Ferroptosis of Microglia in Aging Human White Matter Injury. Ann Neurol. 2023;94(6):1048–1066. doi:10.1002/ana.26770

58. Kenkhuis B, Somarakis A, de Haan L, et al. Iron loading is a prominent feature of activated microglia in Alzheimer’s disease patients. Acta Neuropathol Commun. 2021;9(1):27. doi:10.1186/s40478-021-01126-5

59. Ryan SK, Zelic M, Han Y, et al. Microglia ferroptosis is regulated by SEC24B and contributes to neurodegeneration. Nat Neurosci. 2023;26(1):12–26. doi:10.1038/s41593-022-01221-3

60. Cheli VT, Correale J, Paez PM, Pasquini JM. Iron Metabolism in Oligodendrocytes and Astrocytes, Implications for Myelination and Remyelination. ASN Neuro. 2020;12:175909142096268. doi:10.1177/1759091420962681

61. Absinta M, Sati P, Masuzzo F, et al. Association of Chronic Active Multiple Sclerosis Lesions With Disability In Vivo. JAMA Neurol. 2019;76(12):1474–1483. doi:10.1001/jamaneurol.2019.2399

62. B Y S H, P van G, et al. 7 Tesla magnetic resonance imaging to detect cortical pathology in multiple sclerosis. PloS One. 2014;9(10). doi:10.1371/journal.pone.0108863

