## Supplementary Data for "Aberrant iron deposition in the multiple sclerosis spinal cord relates to neurodegeneration"

#### *Supplementary Data 1* – Diaminobenzidine (DAB)-Enhanced Turnbull Staining Protocol (Tirmann-Schmeltzer Method)

After deparaffinization in xylene and rehydration, sections were washed in deionized water for 5 minutes then immersed in aqueous ammoniumsulfide solution [2% ammonium sulfide in deionized water] for four hours for detection of total non-heme iron then washed with deionized water. Next samples were incubated in a solution of 10% potassium ferricyanide and 1.4% hydrochloric acid in distilled water for 30 minutes at 37°C in the dark. The aforementioned reactions were performed in glass apparatus that was washed with 10% hydrochloric acid and distilled water. After five 3-minutes washing steps with deionized water, sections were incubated in methanol containing 0.3% hydrogen peroxide for 60 minutes. Next, sections were washed with PBS then mounted using Sequenzas. The slides were then developed using DAB (ImmPACT DAB Kit) for 10 minutes, washed 3x with deionized water, and counterstained using hematoxylin.

The figure below shows a thoracic spinal cord section from an MS case with high Turnbull signal in microglia-macrophages (second row), in subpial axons (third row) and in subpial astrocytes (bottom row). DAB-enhanced Turnbull was done in parallel in three adjacent sections which were incubated for 72h in (1) tap water, (2) 0.1M sodium citrate in HCl pH1 buffer, and (3) 0.1M EDTA neutral pH. Turnbull signal was abolished after acid citrate incubation and markedly reduced after incubation with EDTA, although the latter also damaged tissue integrity.

DAB-enhanced Turnbull staining after 72h incubation in: Tap Water or

0.1M Sodium citrate in HCl pH1 buffer

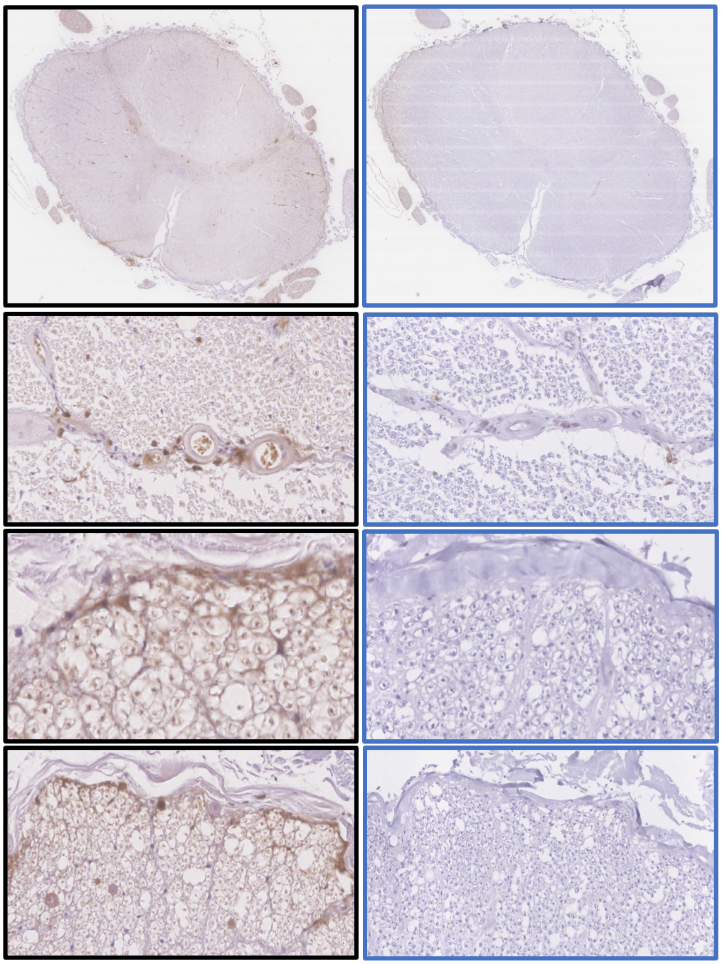

***Supplementary Data 2 - immunohistochemical method***

Formalin-fixed, paraffin-embedded tissue blocks were cut into 6-μm-thick adjacent sections for immunohistochemistry (IHC) using optimised method. Briefly, adjacent sections were baked at 60°C for 20 minutes, deparaffinised in xylene and rehydrated with successive ethanol baths before removal of endogenous peroxidase with 3% $H_2O_2$. Antigen retrieval was optimised for each primary antibody. Adjacent sections were incubated with primary antibodies for: myelin (PLP, BioRad, # MCA839G) and microglia-macrophages (Iba-1, Wako, #019-19741 and CD68, Dako, #M087601-2). Subsequent labelling with secondary antibody and 3,3'-diaminobenzidine (DAB) visualization using the Dako Envision kit. The omission of primary and secondary antibodies separately were used as negative controls. Sections were counter-stained with haematoxylin.

| Antigen | Host | Manufacturer, serial number | Concentration | Incubation | Blocker | Antigen retrieval method |
|---|---|---|---|---|---|---|
| **PLP** | Mouse | Biorad, #MCA839G | 1:1000 | 1h at 20°C | None | Microwave citrate |
| **Iba-1** | Rabbit | Wako, #019-19741 | 1:1000, for double labelling 1:250 | 2h at 20°C | 10% fetal-calf serum | Autoclave citrate |
| **CD68** | Mouse | Dako, #M087601-2 | 1:100 | 1h at 20°C | Serum-free blocker (Dako, #X0909) | Autoclave citrate |
| **Olig-2** | Rabbit | IBL-America, #18953 | 1:100 | Overnight, 4°C | Serum-free blocker (Dako, #X0909) | Autoclave citrate |
| **GFAP** | Rabbit | Dako, #Z0334 | 1:4000 | 1h at 20°C | Serum-free blocker (Dako, #X0909) | Microwave citrate |
| **β-APP** | Mouse | Invitrogen, LN27 #13-0200 | 1:500 | 1h at 20°C | Serum-free blocker (Dako, #X0909) | Autoclave citrate |
| **SMI-32** | Mouse | Merck Millipore, NF-H Non-phosphorylated #Ne1023 | 1:3000 | 1h at 20°C | Serum-free blocker (Dako, #X0909) | Microwave citrate |
| **NPY-Y1R** | Rabbit | Origene, #TA346235 | 1:300 | 1h at 20°C | Serum-free blocker (Dako, #X0909) | Autoclave Tris-EDTA |
| **E06** | Mouse | Absolute, #Ab02746-1.1 | 1:100 | 1h at 20°C | Serum-free blocker (Dako, #X0909) | Autoclave citrate |

### *Supplementary Data 3* – Quantification of microglia-macrophage and axonal injury markers

A subset of cases was selected for additional quantitative analyses of the iron distribution in the microglia-macrophage compartment, and of the association of iron-positive axons with markers of oxidative stress, acute axonal injury, and Wallerian degeneration.

1) For the glial component of the study, a subset of MS cases (n=10) and controls (n=5) propensity-score matched for age at death, sex, and total iron-positive glia number score were stained for: Iba-1, Olig-2, Turnbull & Iba-1 double-labelling, and Turnbull & Olig-2 double-labelling.
2) For analysis of axons, a subset of 5 MS cases with iron-positive axonal clusters, 5 MS cases without iron-positive axonal clusters, and controls (n=5) propensity-score matched for age and sex were stained for: Turnbull, acute axonal injury (β-APP for impaired fast axonal transport, and SMI-32 for non-phosphorylated NF-H), Wallerian degeneration (Neuropeptide-Y-receptor-1 - NPY-Y1R, as previously reported[1]), and oxidative stress (E06 for oxidated phospholipids).

Stained slides were scanned using a Leica Systems Aperio ScanScope AT slide scanner before being imported to QuPath v0.4.3 for quantification. Regions were selected for analysis by placing 40X magnification (361μm x 270μm) fields of view (FOVs) in pre-defined trajectories, selected for quantification in notional tracts. Where possible, according to tissue damage, FOVs were placed in the anterior corticospinal tracts (n=6); lateral corticospinal tracts (n=10); and dorsal columns (n=12), as depicted below. FOVs were not placed in notional tracts containing plaques, which were defined as regions devoid of myelin staining on an adjacent PLP stained section.

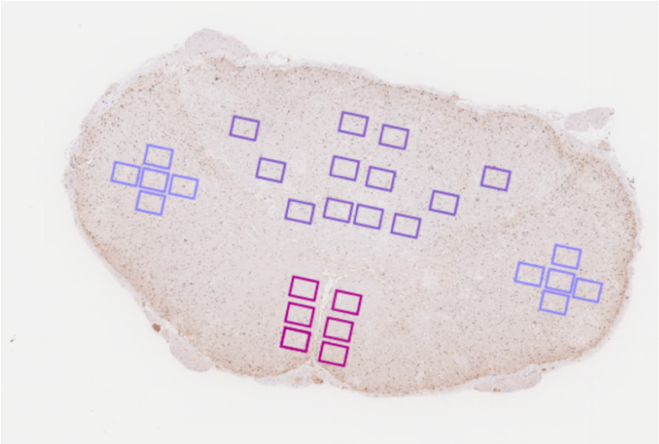

**Supplementary Figure 3: Axial view of MS spinal cord stained for Iba-1 depicting FOV placement in pre-defined trajectories.** FOVs were placed in the anterior corticospinal tract (pink); lateral corticospinal tract (light purple); and dorsal columns (dark purple).

Overall, The number of FOVs placed across all cases for each marker, was as follows: Iba-1-labelled (n=394), Olig-2-labelled (n=394), Iba-1 & Turnbull double-labelled (n=395); Olig-2 & Turnbull double-labelled (n=392), Turnbull-labelled (n=376), E06-labelled (n=380), β-APP-labelled (n=380), SMI-32-labelled (n=380), and NPY-Y1R-labelled (n=378).

In order to quantify total Olig-2-positive cells and E06-, BAPP-, SMI-32- and NPY-1R-positive axons, we used an optimised positive cell/element count macro on QuPath. These macros were verified by blinded manual counts on selected FOVs across all cases (n=15). Test-retest reliability between the optimised macro and manual counting was excellent for Olig-2 (Cronbach's alpha, α=0.997), E06 (Cronbach's alpha, α=0.993); β-APP (Cronbach's alpha, α=0.993); SMI-32 (Cronbach's alpha, α=0.999); and NPY-Y1R (Cronbach's alpha, α=0.995).

To quantify total Iba-1-positive cells, we used a pixel count macro because the staining is non-nuclear. Given that the Turnbull signal was not amenable to automated quantification, Turnbull-positive axons, as well as the number of Iba-1 & Turnbull double-labelled and Olig-2 & Turnbull double-labelled cells was quantified with a blinded manual count, verified by two independent counters.

#### *Supplementary Data 4 – Myelin fraction measurement*

Myelin fraction was calculated as the ratio between the PLP-positive area and the spinal cord section area. To allow the identification of individual myelin wraps in each section, PLP-positive area was measured on QuPath using a very high resolution (0.5 µm/pixel) positive-pixel thresholder (≥0.3) filtering the DAB channel. The picture below shows an example of PLP staining in a non-lesional white matter area; on the right the calculated PLP-positive area is coloured in light blu.

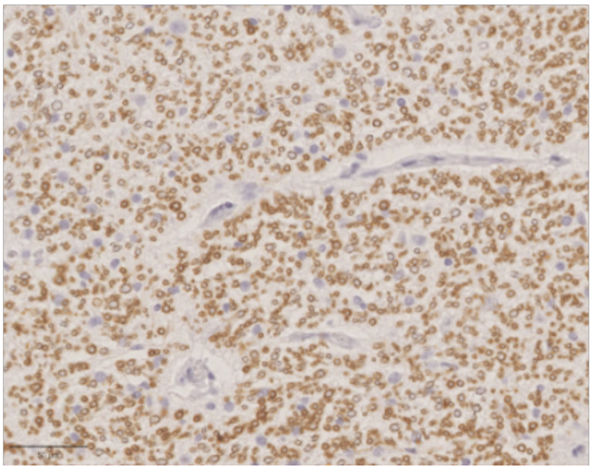

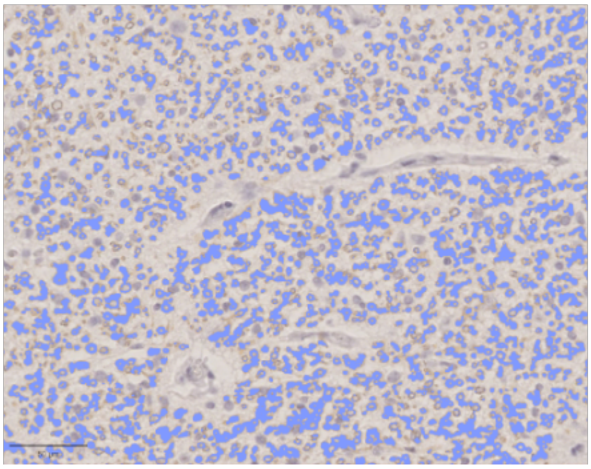

#### *Supplementary Data 5 – Non-heme iron quantification using ICP-OES*

Two 15µm thick sections were taken for each FFPE spinal cord sample. Tissue was baked at 60'C for 40' and incubated in xylene for 15 minutes for deparaffinization. After two washes with 70% and 90% alcohol, sections were incubated for 72h in 1mL of 0.1M sodium citrate pH 1.0 solution. The 1 mL buffer was extracted and sent for quantification with inductively coupled plasma optical emission spectroscopy (ICP-OES).

The inductively coupled plasma optical emission spectroscopy (ICP-OES) analysis was carried out on a sequential ICP (Ultima, Horiba Jobin Yvon, France), using the Horiba Jobin Yvon ICP Analyst software (v. 5.4). A monochromator with a Czerny-Turner spectrometer was used. The plasma in the ICP-OES was made by partially ionizing argon gas. The instrument configuration and experimental conditions of the ICP-OES analysis are summarized below:

**Analytical conditions for ICP-OES analysis.**

| Instrument | ICP Ultima |
|---|---|
| RF generator power | 1.05 kW |
| RF frequency | 40.68 MHz |
| Plasma gas flow rate | 12 L $min^{-1}$ |
| Carrier gas flow rate | 1.0 L $min^{-1}$ |
| Sample introduction | Miramist nebulizer |
| Misting chamber | Cyclonic glass chamber |
| Observation method | Radial |
| Injector tube diameter | 3 mm |

Iron content was determined from emission readings at the Fe 259.940 nm emission line. The instrument was calibrated with dilutions with buffer solution of an iron standard (ICP standard, 1000 $mgL^{-1}$).

***Supplementary Data 6 - Iron-positive axons associate with greater oxidative stress but not with markers of acute axonal injury***

Iron-positive axons were manually counted in in a subset of MS cases and controls in predefined trajectories in the anterior, lateral and posterior funiculi.

For this purpose, a subset of 10 MS cases and 5 controls were propensity score-matched for sex and age at death; MS cases were selected to include 5 cases with iron-positive axons (i.e. semiquantitative score for iron-positive axons > 0) and 5 cases without iron-positive axons (i.e. semiquantitative score for iron-positive axons = 0).

| | **MS (n=10)** | **Controls (n=5)** |
|---|---|---|
| **Sex** | Male=3; Female=7 | Male=2; Female=3 |
| **Age at death (years)** | 59.4 (40-75) | 70.4 (64-80) |
| **Brain weight (grams)** | 1216.1 (973-1380) | 1309 (1130-1443) |
| **Post-mortem interval (hours)** | 15.5 (6-27) | 37.2 (23-52) |
| **Cerebrospinal fluid pH** | 7.26 (6.2-9.2) | 7.75 (7.7-7.8) |
| **Disease duration (years)** | 29.1 (15-58) | N/A |
| **Clinical MS classification** | PPMS=2; SPMS=7; N/A=1 | N/A |

| | | |
|---|---|---|
| **Time to wheelchair (years)** | 18.8 (6-50) | N/A |

**Supplementary Table 6: Characteristics of MS cases and controls comprising the axonal subset.** Mean values with the range in parentheses are provided where applicable. N/A, not applicable or not available.

A 40-fold increase in the number of iron-positive axons was observed in MS compared to controls (A). The presence of iron-positive axons in a funiculus associated with greater number of manually counted axons positive for oxidative stress (oxidated phospholipids, E06; B). No association was seen instead with the number of axons positive for acute axonal injury (beta-amyloid precursor protein – B-APP positive axonal spheroids; C), axonal damage (non-phosphorylated neurofilament H - SMI-32; D) or Wallerian degeneration (Neuropeptide Y receptor 1 – NPY1R; E). Pairwise comparison corrected for case-effect using GEE model, ** indicates p<0.01.

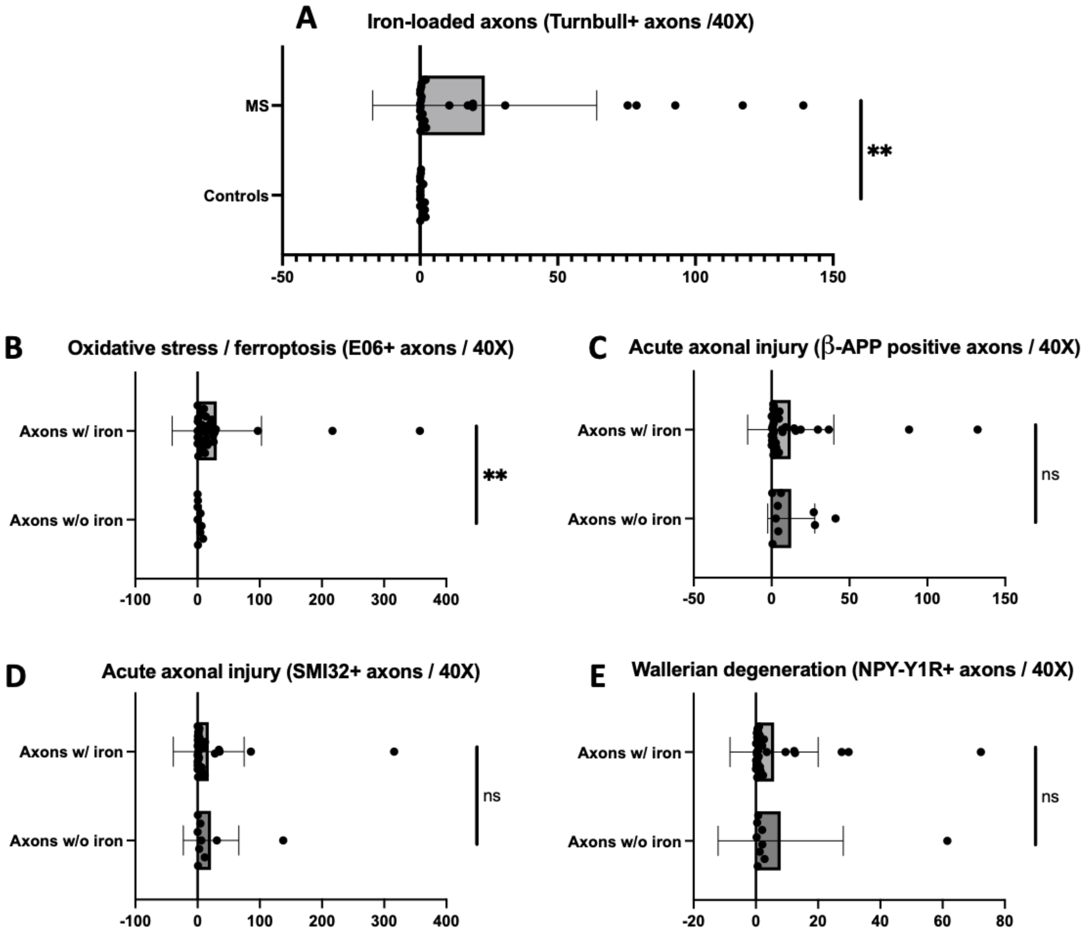


***Supplementary Data 7 – Iron-positive microglia-macrophages are selectively increased in MS, with no difference in iron-positive oligodendrocytes.***

Iron distribution in glial cells was quantified in a subset of MS cases (n=10) and controls (n=5) in predefined trajectories in the anterior, lateral and posterior funiculi.

For this purpose, a subset of MS cases (n=10) and controls (n=5) were propensity-score matched for sex, age at death, and the total glia semiquantitative score.

| | **MS (n=10)** | **Controls (n=5)** |
|---|---|---|
| **Sex** | Male=5; Female=5 | Male=3; Female=2 |
| **Age at death (years)** | 73.1 (48-92) | 76.8 (68-85) |
| **Brain weight (grams)** | 1165.4 (982-1366) | 1218 (1108-1394) |
| **Post-mortem interval (hours)** | 16.6 (8-22) | 26 (21-33) |
| **Cerebrospinal fluid pH** | 6.87 (6.45-7.25) | 7.625 (7.2-7.8) |
| **Disease duration (years)** | 38.3 (26-58) | N/A |
| **Clinical MS classification** | PPMS=1; SPMS=8; N/A=1 | N/A |
| **Time to wheelchair (years)** | 24.1 (0-50) | N/A |

**Supplementary Table 7: Characteristics of MS cases and controls comprising the glial subset.** Mean values with the range in parentheses are provided where applicable. N/A, not applicable.

We observed no difference in the number of Olig-2 cells (total oligodendrocytes, A), Olig-2 & Turnbull-positive cells (i.e. iron-positive oligodendrocytes, B), and in the proportion of Olig-2 & Turnbull-positive cells on total Olig-2 cells (C) between MS cases and controls.

No significant difference was found between the average number of Iba-1 cells in MS cases and controls (A). We did, however, observe a significant, near seven-fold increase in the average number of Iba-1 double-positive cells in MS cases compared to controls (Wald chi-square, $\chi^2$=6.009, p=0.014, B). When Iba-1 double-positive cells were considered as a proportion of Iba-1 single-positive cells, the same result was found (Wald chi-square, $\chi^2$=5.800, p=0.016, C).
No difference was found after correcting for age.

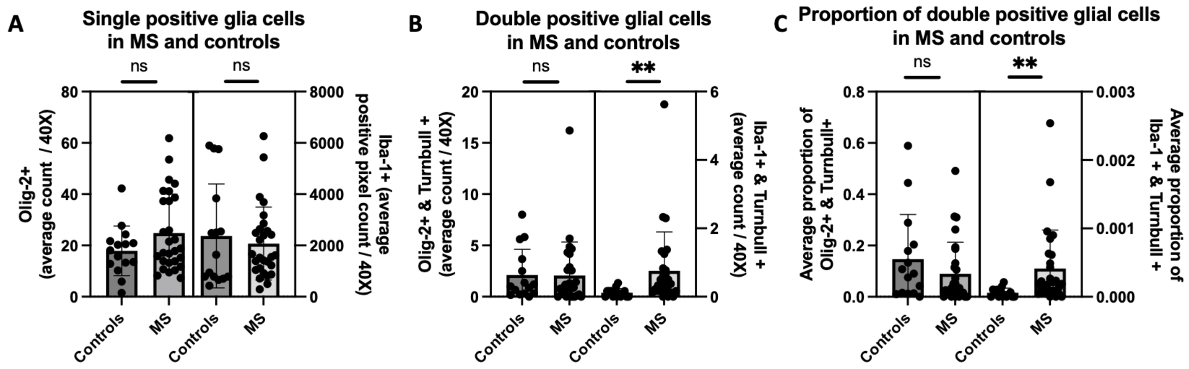
A
Single positive glia cells in MS and controls
ns
ns
Olig-2+ (average count / 40X)
Iba-1+ (average positive pixel count / 40X)
0
20
40
60
80
0
2000
4000
6000
8000
Controls
MS
Controls
MS
B
Double positive glial cells in MS and controls
ns
**
Olig-2+ & Turnbull + (average count / 40X)
Iba-1+ & Turnbull + (average count / 40X)
0
5
10
15
20
0
2
4
6
Controls
MS
Controls
MS
C
Proportion of double positive glial cells in MS and controls
ns
**
Average proportion of Olig-2+ & Turnbull+
Average proportion of Iba-1 + & Turnbull +
0.0
0.2
0.4
0.6
0.8
0.000
0.001
0.002
0.003
Controls
MS
Controls
MS
